# Joint inference of migration and reassortment patterns for viruses with segment genomes

**DOI:** 10.1101/2021.05.15.442587

**Authors:** Ugnė Stolz, Tanja Stadler, Nicola F. Müller, Timothy G. Vaughan

## Abstract

The structured coalescent allows inferring migration patterns between viral sub-populations from genetic sequence data. However, these analyses typically assume that no genetic recombination process impacted the sequence evolution of pathogens. For segmented viruses, such as influenza, that can undergo reassortment this assumption is broken. Reassortment reshuffles the segments of different parent lineages upon a coinfection event, which means that the shared history of viruses has to be represented by a network instead of a tree. Therefore, full genome analyses of such viruses is complex or even impossible. While this problem has been addressed for unstructured populations, it is still impossible to account for population structure, such as induced by different host populations, while also accounting for reassortmentWe address this by extending the structured coalescent to account for reassortment and present a framework for investigating possible ties between reassortment and migration (host jump) events. This method can accurately estimate sub-population dependent effective populations sizes, reassortment and migration rates from simulated data. Additionally, we apply the new model to avian influenza A/H5N1 sequences, sampled from two avian host types, Anseriformes and Galliformes. We contrast our results with a structured coalescent without reassortment inference, which assumes independently evolving segments. This reveals that taking into account segment reassortment and using sequencing data from several viral segments for joint phylodynamic inference leads to different estimates for effective population sizes, migration and clock rates.

This new model is implemented as the Structured Coalescent with Reassortment (SCoRe) package for BEAST 2.5 and is available at https://github.com/jugne/SCORE.

## Introduction

Influenza viruses are continuously evolving, escaping host immunity or switching host species. Additionally, influenza diversity is promoted by intermediate mammalian hosts, such as swine, which can serve as a mixing vessel for human, avian and its own influenza strains, with subsequent spill over back to the human population (Ma, Kahn, and Richt 2009; Khiabanian, Trifonov, and Rabadan 2009). This pattern usually manifests through *reassortment* - a form of recombination in segmented viruses - when divergent strains infect a single host cell and exchange genetic segments upon replication (McDonald et al. 2016).

Like other forms of recombination, reassortment poses challenges for phylodynamic inference methods, which use influenza sequences to gain insights into the transmission history of samples and values of epidemiological parameters. This is because, without an explicit model for the reassortment process, genomic segments must be analyzed in isolation in order to avoid making incorrect assumptions about the degree to which ancestry is shared between segments.

For this reason, we recently introduced the coalescent with reassortment model (motivated by the coalescent with recombination model of Hudson (1983)) which explicitly accounts for reassortment between influenza genome segments (Müller, Stolz, et al. 2020). Together with a Markov chain Monte Carlo (MCMC) algorithm for sampling from the posterior distribution of reassortment networks, this allows genetic data from all genome segments to be incorporated into a single Bayesian phylodynamic analysis, even in the presence of substantial reassortment.

Here, we extend this model in order to account for population structure, where sub-populations (types) can be associated with discrete-valued lineage traits such as the host species or coarse-grained geographic location. We accomplish this by building on previous work (Müller, Rasmussen, and Stadler 2017; Müller, Rasmussen, and Stadler 2018) which allows for relatively efficient inference by explicitly marginalizing over all migration histories. We develop both exact and approximate approaches for sampling from the posterior distribution of reassortment networks, reassortment rates, effective population sizes and migration rates under the combined model: the structured coalescent with reassortment (SCoRe). In order to gain information about when and where migration events occurred, we also implement a stochastic mapping technique (Nielsen 2002; Huelsenbeck, Nielsen, and Bollback 2003) that allows us to sample migration histories on the reassortment networks.

We demonstrate that SCoRe is able to correctly estimate the effective population sizes, reassortment and migration rates when applied to simulated data. We then show that assuming independently evolving viral segments can bias these inferences. Next, we apply the structured coalescent with reassortment to an avian influenza A/H5N1 dataset with isolates from two bird orders: Anseriformes and Galliformes. Anseriformes are waterfowl birds that have been previously identified as a reservoir for avian influenza viruses (Webster et al. 2006; Kim et al. 2009). Migratory patterns of wild Anseriformes facilitate the spread of influenza between different locations, while domesticated Anseriformes (primarily ducks) develop less pathogenic infection and play a significant role in transmitting to other domestic poultry (Hulse-Post et al. 2005; Li et al. 2004). Such spill-over from Anseriformes to Galliformes (ground-feeding, mostly domesticated birds) in turn can cause further transmission to non-avian livestock as well as humans. Our analysis is able to recover this transmission pattern and shows different estimates of model parameters than the ones obtained when assuming independently evolving segments. We also investigate possible correlation between reassortment and migration (host jump) events, setting a framework for such studies on different influenza strains and host types.

### New Approaches

Previously, performing model-based Bayesian phylogeographic inference of the migration patterns of segmented viruses required either a) assuming that reassortment is frequent and thus that segments evolved according to independent phylogenies, b) assuming that reassortment is rare and thus that all segments share a single phylogeny, or c) limiting the analysis to a single segment. The first two possibilities make assumptions that lead to bias when not met, while the third is wasteful in that it involves throwing away potentially-useful information. We present here the structured coalescent with reassortment model, which provides the basis for an alternative approach that allows all segments to be analyzed together. This model is an extension of the structured coalescent, and considers the tree to be the outcome of a backward-time process where lineages can coalesce and reassort within, and migrate between, sub-populations.

To obtain a practical inference method based on this model, we extend the marginal approximation of the structured coalescent (Müller, Rasmussen, and Stadler 2017; Müller, Rasmussen, and Stadler 2018) to account for reassortment events. This involves numerically solving a set of differential equations describing the change in ancestral lineage state probabilities over time. Combined with our recently-developed MCMC algorithm for sampling reassortment networks(Müller, Stolz, et al. 2020), yields our approach for sampling from the joint posterior distribution of these networks, the embedding of segments trees within those networks, migration histories, and other associated parameters.

Additionally, we have developed a stochastic mapping algorithm to impute individual migration events over the network. Once this mapping is complete, each network sampled from the posterior distribution is annotated with a possible sequence of reassortment, coalescent and migration events and can therefore be used to investigate possible relationships and correlations between these events.

An early version of this method was part of the Masters thesis of the first author (Jankauskaite 2019).

## Results

### Implementation validation and true network parameter estimation

We first ensure that our implementation of the exact variant of SCoRe (SCoRe-exact) samples from the true distribution of structured coalescent with reassortment. Additionally, we show that the implementation of the approximate version of SCoRe does not qualitatively distort shapes and summary statistics of these distributions. To do this, we used our MCMC algorithm to produce ensembles of reassortment networks under (a) SCoRe-exact and (b) SCoRe. We then used direct simulation (Gillespi 1976) to produce a third set of networks simulated under the same set of parameter values. Next, we compared the frequency distributions of network height, length, and reassortment node count from each of these ensembles (supplementary figures S16, S17). To measure the difference of distributions more precisely, we calculated the Kolmogorov-Smirnov (KS) statistic as a function of iteration count. In the exact case, the KS differences asymptote to around 0.01 − 0.003, and from this we conclude that distributions of network statistics match with high precision. For the approximation, the KS differences are slightly larger (< 0.1), but the distributions maintain similar shapes and mean values. For the analyses below, we only apply the approximate version of SCoRe, as it is substantially faster allowing us to apply it to considerably larger datasets (see Materials and Methods).

To demonstrate that SCoRe allows us to estimate the model parameters correctly, we considered two different well-calibrated simulation studies. First, we test how well SCoRe is able to recover effective population sizes, reassortment and migration rates when the true network is known and fixed (supplementary figure S11). We then test the ability of SCoRe to jointly infer the network and its parameters was studied for high, low and mixed segment clock rates and two different migration priors (see Materials and Methods). Figure 1 shows the results for high (5 × 10^−3^) clock rates and exponential migration prior. Supplementary figures S12-S14 show the results for remaining combinations of migration rate priors and clock rates. Overall, between 91 and 98 per cent of true parameter values fell within the 95% HPD interval (supplementary table S1.

**Figure 1:**
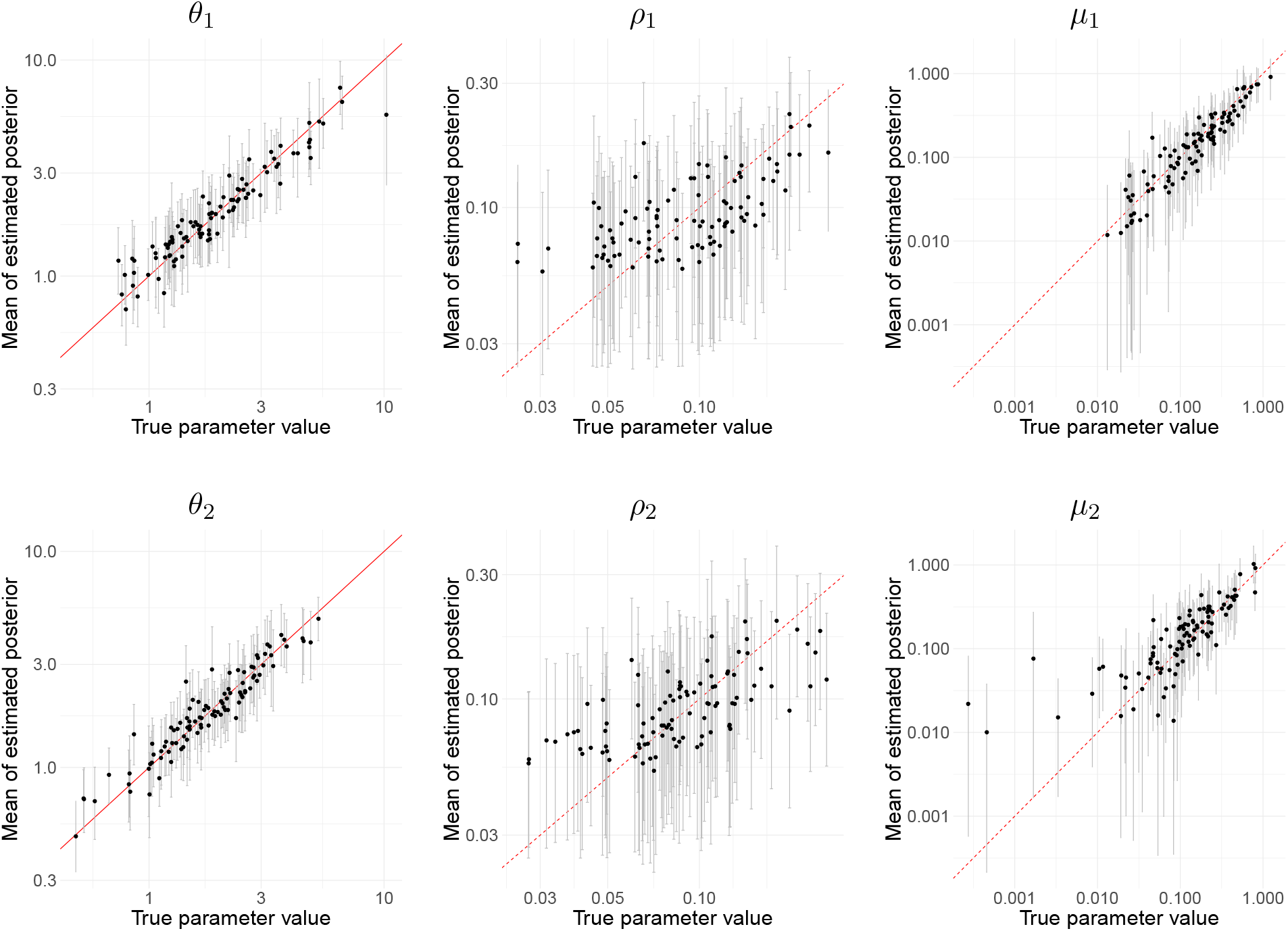
Inference of effective population size (*θ*), reassortment (*ρ*) and migration (*µ*) rates from 100 simulated genetic sequence data for 2 types with exponential migration rate prior and high clock rate (5 *×* 10^*-*3^ for all 4 segments). First row of is for type 1 and second – for type 2 parameters. True (x-axis) versus estimated (y-axis) effective population sizes. Grey bars are 95% confidence intervals, red marks the x=y curve.

### Joint inference from viral segments reduces the relative error of model parameters

In our final simulation study, we compared the relative error of effective population size and migration rate posterior distributions inferred under SCoRe when the true reassortment rate is known to the distributions obtained assuming independent genomic segments under the structured coalescent model that does not include reassortment (MASCOT, Müller, Rasmussen, and Stadler 2018). The true effective population sizes, migration and reassortment rates were randomly drawn from their respective known prior distributions. We obtained 100 such sets of parameter values and simulated sequences for 4 segments and 2 sub-populations repeatedly with low (5 × 10^−4^) and high (5 × 10^−3^) clock rates (see Materials and Methods for more details).

Both inference methods were supplied with simulated sequences. The prior distributions for effective population sizes and migration rates were set to those from which true values were drawn in the simulation. Additionally, reassortment rates were not estimated and set to the true values in SCoRe. Both methods had similar accuracy for low clock rates, with median relative error being slightly smaller for SCoRe compared to MASCOT. In the case of high clock rates, the difference was more pronounced with median relative error being up to 5.4% smaller for migration rates for SCoRe compared to MASCOT. (supplementary figure S15).

### Network and its parameter inference for avian influenza A/H5N1

We next assembled genetic sequences of influenza A/H5N1 viruses sampled between 2008 and 2016 and grouped them into Anseriformes or Galliformes, according to the order of the host species of the isolate (see Materials and Methods). For each of those samples we used two segments, the two surface proteins HA and NA, for further analysis. We then randomly subsampled this dataset into 10 smaller subsets, each containing 200 sequences and ran three independent analyses for each subset under SCoRe and MASCOT in BEAST 2.5 (R. Bouckaert et al. 2018), using parallel tempering (Altekar et al. 2004; Müller and R. R. Bouckaert 2020). On each segment, we allowed for different evolutionary rates of the first two and the third codon position evolving under an HKY+Γ_4_ (Hasegawa, Kishino, and Yano 1985; Yang 1994) substitution model.

The inferred posterior distributions for migration rates are bimodal (figure 2, supplementary figures S1, S2, “Unfiltered”) and the inferred root bird order varies with the two different modes assignment of the segment trees (MASCOT) and the network (SCoRe). Furthermore, maximum clade credibility (MCC) and maximum posterior networks show that the inferred root state strongly correlates with the majority of the sampled A/H5N1 evolution occurring in the same state (supplementary figures S4, S5). Ewing and Nicholls 2004 discuss such bimodality in migration rates for MCMC structured tree inference methods and suggest using additional prior knowledge to weight one of the migration directions. In this case it has been shown that migratory water-fowl birds (Anseriformes) are the main natural influenza A/H5N1 reservoir (Olsen et al. 2006; Webster et al. 2006; Kim et al. 2009; Trovão et al. 2015). Furthermore, H5N1 is considered to be less pathogenic to domesticated ducks (also Anseriformes) than other poultry thus prolonging the disease shedding period and promoting introduction into the domesticated bird populations (Hulse-Post et al. 2005; Li et al. 2004). Therefore, to obtain the conditioned posterior distribution, we decided to condition on the root node type. We filter the ensembles produced by MCMC, retaining only those samples in which both segment roots are associated with Anseriformes (see Methods and Methods). Figure 2 and supplementary figures S1, S2 show that conditioning completely removed bimodality of estimated migration rate distributions for most subsets with SCoRe and all subsets with MASCOT. We rely on filtered posterior distributions in the further analyses and present equivalent figures obtained before filtering for consistency.

**Figure 2:**
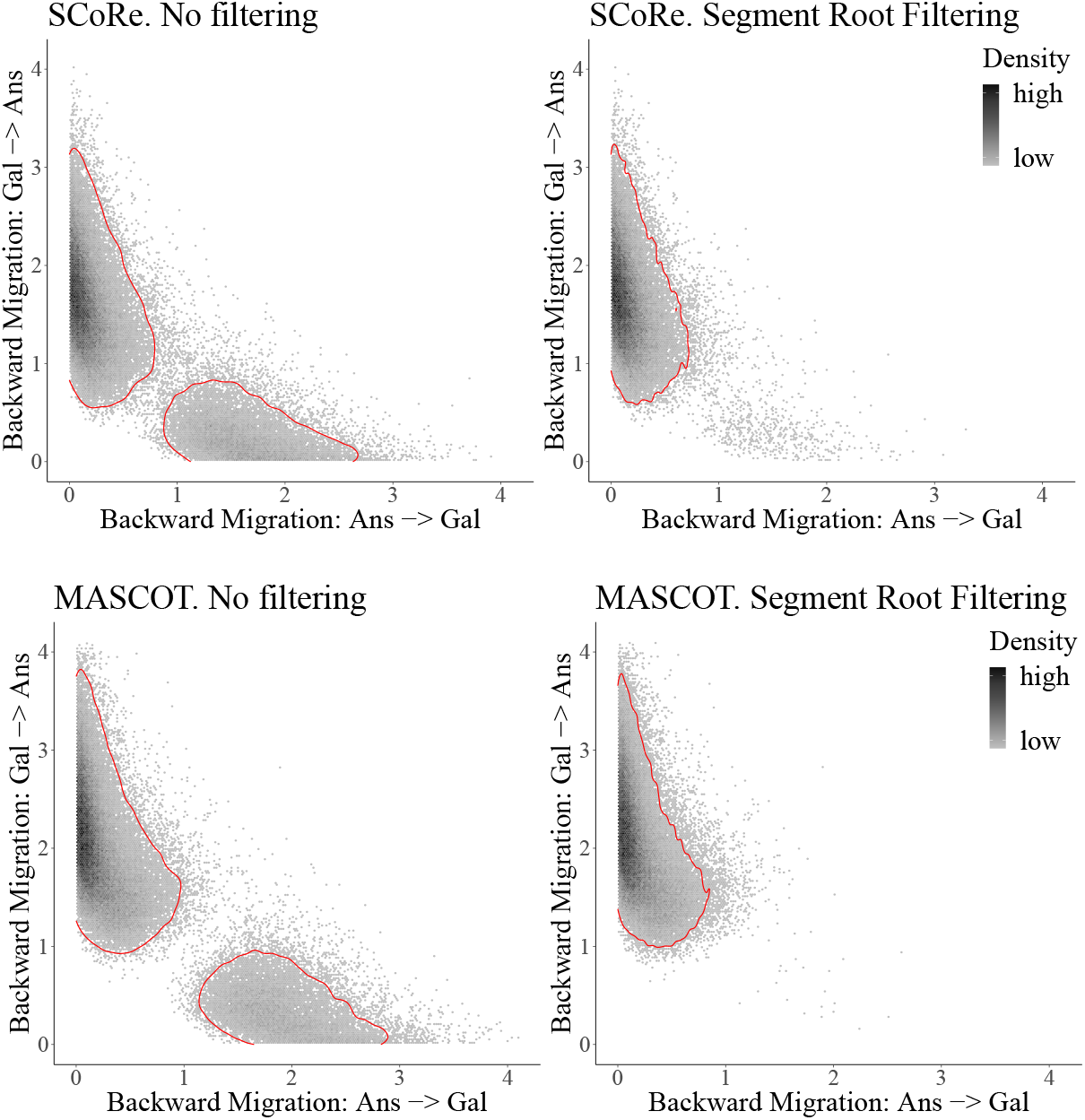
2D density of backward in time migration rate posterior estimates for all 10 subsets before and after segment root filtering for MASCOT and SCoRe. Red contour line marks the 95% HPD area for the estimated 2D density.

With this condition in place, both SCoRe and MASCOT recover the expected trajectory of the virus, with high backwards in time migration from ground feeding birds (Galliformes) to water-fowl (Anseriformes) (see figure S3). Backwards migration rate from Anseriformes to Galliformes is, in comparison, minuscule. The root of MCC network as well as majority of the network length is in the Anseriformes for all 10 subsets (figure 3, supplementary figure S6).

**Figure 3:**
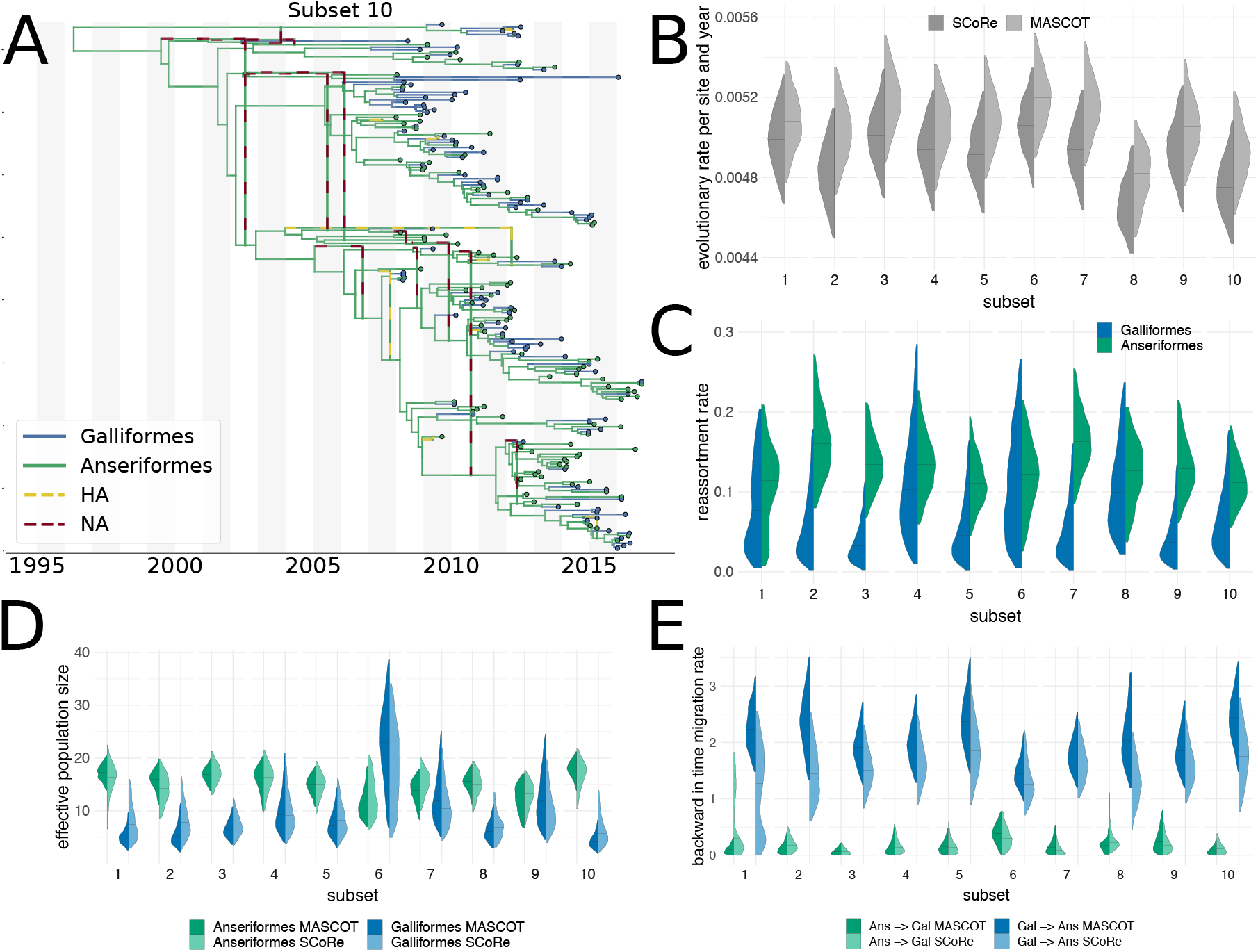
MCC network and posterior parameter estimates. Segment root type filtering applied. The distributions are compared for both host types (Anseriformes and Galliformes) and for MASCOT and SCoRe packages. **A** An example maximum clade credibility network for random subset 10 of A/H5N1. Dashed lines show segment movement at reassortment events. **B** Comparison of inferred clock rates. SCoRe obtains lower estimates than MASCOT. **C** Comparison of reassortment rates inferred by SCoRe for two bird orders. **D** Comparison of effective population sizes. **E** Comparison of inferred backwards in time migration rates for Anseriformes (Ans) and Galliformes (Gal).

### Correlation between reassortment and migration events in avian influenza A/H5N1

Next, we test for the presence of an association between reassortment events and host jumps (migration events). To do so, we define a short time window immediately before each host jump event in the network. We then compute the empirical rate of reassortment events within these windows (“on window”) and compare these to the corresponding rates for all parts of the network that fall outside of these windows (“off window”). We compute these rates for all networks sampled from the posterior distribution of reassortment networks and then compute the difference between the rates within and outside of the window. If this difference is above zero, the rate of reassortment is higher within the windows, i.e. before host jump events. If the difference is below zero, this suggests lower rates of reassortment before host jump events.

Additionally, we compute the same difference for networks simulated under the structured coalescent with reassortment and the inferred parameters, that is for posterior predictive simulations. Since a fitness effect of reassortment was shown previously from human influenza viuses (Müller, Stolz, et al. 2020), we also tried to disentangle general fitness benefits of reassortment from its association with host jumps. To do so, we split all networks into “fit” and “unfit” based on how long the descendants of an edge persist into the future. If they persist for more than two years, we classify an edge as fit and as unfit otherwise. We then again compute the same difference between reassortment rates on and off windows for only for edges either classified as “fit” or “unfit”.

As shown in figure 4, we see a slight increase in reassortment rates before host hump events for all 10 subsets. In all cases, however, the 95% Highest posterior density interval still includes no increase in rates of reassortment before host jump events. For most subsets, we also find a slight increase in rates of reassortment before host jump events when looking only at fit respectively unfit parts of the networks. These estimates, however, are very uncertain and still all include no association at all, as well. Additionally, we looked at what happens when we change the size of the window around host jump events, as well as the definition of what constitutes a fit and unfit edge (see supplementary figures S7-S10 for different window sizes and fitness distances), which shows largely consistent results.

**Figure 4:**
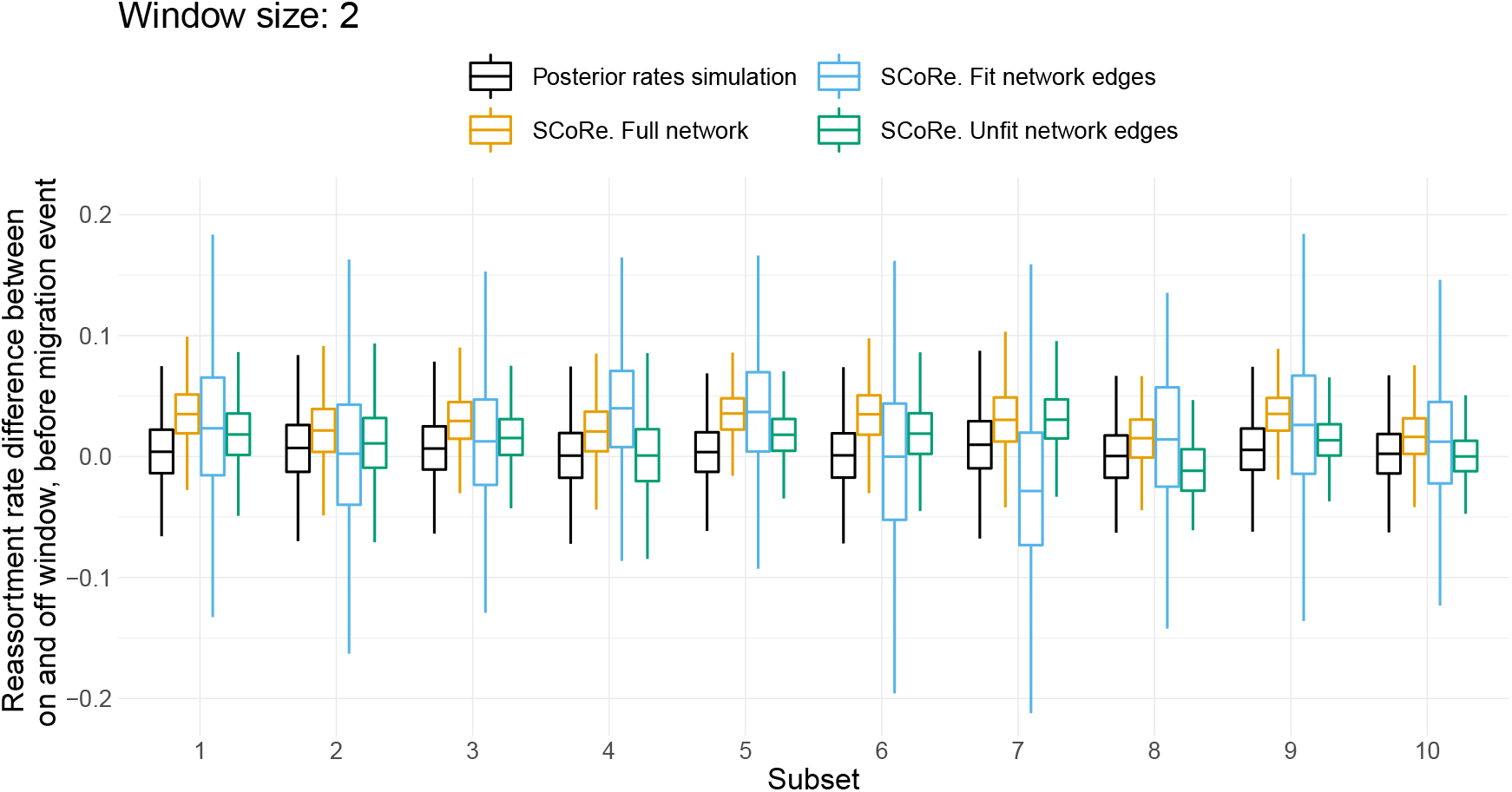
Reassortment rates near host jump events compared to elsewhere. Window size: 2 years. Lineage is fit if it has descendants 2 year or more in the future. For each run, we compare distributions when (a) simulating (no sequencing data) with the posterior rates obtained by the inference or (b) inferring from the sequencing data under the SCoRe model. Since there is a previously defined increase in reassortment due to fitness, which is informed by the data, we show the reassortment rate difference for full network and separately for fit and unfit network edges.

## Discussion

The approach presented here permits inference of reassortment networks from sequences of viruses with segmented genomes while accounting for population structure. This is done by expanding the efficient structured coalescent approach described in Müller, Rasmussen, and Stadler 2017 to the structured coalescent with reassortment. Using several simulation studies, we have shown that, even though approximate, our method can reliably recover effective population sizes, reassortment and migration rates.

When comparing the inference results between when accounting for reassortment compared to assuming segments to evolve independently, we find that both approaches recover similar transmission dynamics for an avian influenza A/H5N1 dataset with two types. The assumption of independent segments, however, leads to quantitatively different estimates for the evolutionary rate and backwards migration rates than those obtained by SCoRe. This is consistent with previous findings for the unstructured case (Müller, Stolz, et al. 2020) where assuming independently evolving segments also lead to higher evolutionary rate estimates. Both models showed a strong bimodality in estimated migration rates. This may be due to insufficient data, the omission of intermediate host types from the model or falsely assuming constant effective population sizes and rates of migration. The first two possibilities could be addressed by using a larger amount of sequences or distinguishing between more host types and or geographic locations. The latter one requires extensions to the model, for example, by allowing for effective population sizes to change over time (Drummond et al. 2005).

We have also investigated possible associations between reassortment and host jump events using a dataset of Haemagglutinin and Neuraminidase avian influenza A/H5N1 sequences. While we find some evidence that reassortment rates are elevated prior to host jump events, the confidence intervals did not exclude no elevated rates. Using a dataset that spans a longer time window could potentially help increase the precision of these estimates. Additionally, the role of reassortment in host switching may be more pronounced for host switches between more distantly related species. Therefore, while we do not find strong evidence for the role of reassortment in host switching in the dataset analysed, we show a potential framework to investigate the role of reassortment in host switching.

## Materials and Methods

### The structured coalescent with reassortment

Here, we extend the coalescent with reassortment (Müller, Stolz, et al. 2020) process to allow for population structure. Each lineage *L* of a network *G*, described by this process, carries a full set of genomic segments, subset of which 𝒞(*L*) are ancestral to the samples. In addition, each network lineage is in a particular type (member of a sub-population) and migration between types is allowed. At any given time, we define a lineage *L* by its type *l*, which takes values from a types set {1, 2, …, *m*} and the ancestral segments 𝒞 (*L*) it carries: *L* = [*l*, 𝒞 (*L*)]. Given *n* coexisting lineages and *m* types, there are *m*^*n*^ possible network configurations 𝒦 := {*L*_*i*_ = [*l*_*i*_, 𝒞 (*L*_*i*_)] | *i ∈* {1, 2, …, *n*}}, w.r.t. the type of each lineage. We model the generation of structured coalescent with reassortment networks as a continuous backward in time Markov chain (CTMC), that generates the network configuration 𝒦 (type in the Markov chain) by *coalescent, migration, reassortment* and *sampling* events.

Under a standard coalescent model, the probability per unit of time that two coexisting lineages have a common ancestor is equal to an inverse of effective population size *Ne*. We only permit coalescent events between any two lineages *L* and *L*′ that are in the same type *a*. Therefore, the pairwise coalescent rate of type *a, λ*_*a*_, is an inverse of sub-population effective population size *Ne*_*a*_. Immediately after a coalescent event, the parent lineage *L*_*p*_ carry ancestral segments of both child lineages (Müller, Stolz, et al. 2020) and inherits their type *L*_*p*_ = [*a*, 𝒞 (*L*) ∪ 𝒞*L*′)]. If *k*_*a*_(𝒦) is the number of lineages in type *a* for some configuration 𝒦, then the total coalescent rate is the sum over all possible types:

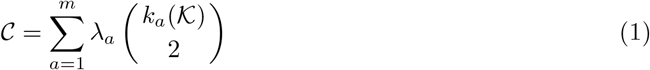

Migration event involves a single lineage and change its type, but not the segments it carries. That is, a migration event from type *a* to *b* on a lineage *L* would change it from *L* = [*a*, 𝒞 (*L*)] to *L*′= [*b*, 𝒞 (*L*)]. Denote the rate of such migration event *µ*_*ab*_ and let *µ*_*aa*_ = 0. Then, the total migration rate for some configuration *K* is the sum over all *n* lineages to change their current type *l*_*i*_ into any other:

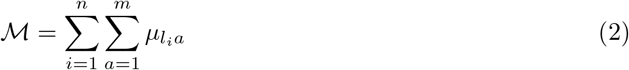

At a reassortment event, we observe a lineage *L* and its two parents *L*_*p*1_ and *L*_*p*2_. Each ancestral segment is inherited by the child lineage from parent *L*_*p*1_ or *L*_*p*2_ with probability *p*. Here, we assume that this probability is equal for both parents: *p* = 0.5. A reassortment event is observable, if at least one one ancestral segment originated from a different parent than all other segments: 𝒞 (*L*) ∩ 𝒞(*L*_*p*1_) ≠ ∅, 𝒞(*L*) ∩ 𝒞(*L*_*p*2_) ≠ ∅ and 𝒞(*L*_*p*1_) ∩ 𝒞(*L*_*p*2_) = ∅. This occurs at a probability (1 − *p*^|*C*(*L*)|−1^), where |𝒞(*L*)| is the number of ancestral segments carried by lineage *L*. Now, let *ρ*_*a*_ be the reassortment rate at any lineage in type *a* and *P*_*t*_(*L*_*j*_ | 𝒦, *G*) an indicator probability of lineage *L*_*j*_ being in the type *a* at time *t*, given the configuration 𝒦 and network *G*. Then, the total rate of observed reassortment is

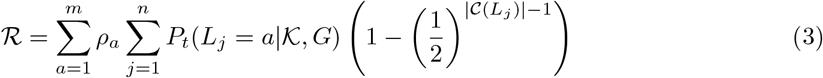

As is standard in coalescent models, we condition on sampling events.

### Target posterior probability

In order to perform MCMC sampling, we describe the target posterior distribution of networks using Bayes theorem.

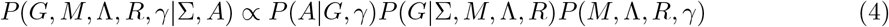

Here, *A* is the set of multiple sequence alignments for each segment and Σ is the types of each sample. *M*, Λ and *R* are respectively sets of migration, coalescent and reassortment rates of the network. Finally, *γ* is a set of substitution model parameters.

The probability of sequence data given the network and substitution rate parameters *P* (*A*|*G, γ*) can be factored into a sum of probabilities given the segments trees and calculated by the Felsenstein pruning algorithm (Felsenstein 1981; Müller, Stolz, et al. 2020). We set the joint probability of network parameters *P* (*M*, Λ, *R, γ*) to be equal to the multiplication of their independent prior probabilities. Next, we explain how to obtain the probability of the structured coalescent with reassortment network *P* (*G*|Σ, *M*, Λ, *R*) and use MASCO approximation (Müller, Rasmussen, and Stadler 2018) to make its calculation feasible.

### The structured coalescent with reassortment as a network prior

As described by Müller, Rasmussen, and Stadler 2017, we seek to marginalise over all possible migration histories *H* to obtain the probability of a network *G*.

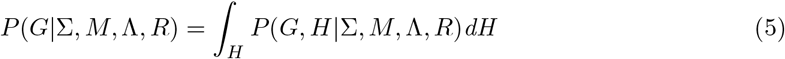

Equivalent to the tree case (Müller, Rasmussen, and Stadler 2017), the above equation for network can be evaluated by

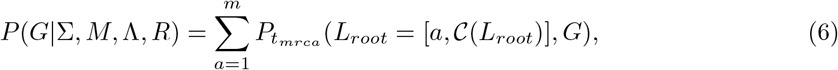

where *P*_*t*_(𝒦, *G*) is a joint probability of the network *G* before the time *t* and the network configuration 𝒦 at this time. When the time *t*_*mrca*_ of the network root is reached, there is only one lineage and the probability reduces to 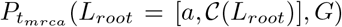. The ancestral segment set of the root lineage 𝒞 (*L*_*root*_) will always contain all segments ancestral to the samples.

### Approximation of the structured coalescent with reassortment

To obtain 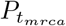we numerically integrate *P*_*t*_ until *t* = *t*_*mrca*_, where *t* = 0 is the time of the first sample and time increases in the past. In turn, *P*_*t*_ can be factored into the contribution of network events (coalescent and reassortment) and the corresponding intervals between these events. Note, that there is no contribution of migration events since we marginalize over all migration histories. We have derived the exact equations for *P*_*t*_(𝒦, *G*) in terms of network events and intervals contributions (see Supplement). However, in order to evaluate them and account for all possible network type configurations 𝒦, we need to numerically solve *m*^*n*^ differential equations, which becomes intractable for larger or highly structured datasets. As in (Müller, Rasmussen, and Stadler 2018), we assume that the state lineages is pairwise independent, give the tree.

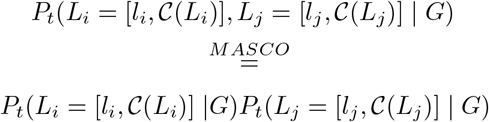

This approximation eliminates the need to jointly account for all possible lineage type configurations. Thus it reduces the number of equations from *m*^*n*^ to *m* × *n*. Next, we show how to extend this approach in the case of structured coalescent with reassortment network.

The previously derived ODEs needed to calculate structured coalescent tree prior (Müller, Rasmussen, and Stadler 2017; Müller, Rasmussen, and Stadler 2018) involves migration and coalescent terms that are equivalent in the structured network case. For completeness, we restate them and include the necessary reassortment terms in order to obtain the approximate structured network prior. To evaluate the change in a marginal lineage type probability *P*_*t*_(*L*_*i*_ = [*l*_*i*_, 𝒞 (*L*_*i*_)], *G*) within a network interval, we obtain its derivative over time. Furthermore, to avoid numerical instability caused by vanishing values (Müller, Rasmussen, and Stadler 2018) we calculate instead *P*_*t*_(*L*_*i*_ = [*l*_*i*_, 𝒞(*L*_*i*_)]|*G*) = *P*_*t*_(*L*_*i*_ = [*l*_*i*_, 𝒞(*L*_*i*_)], *G*)*/P*_*t*_(*G*). Derivative of which is (see Supplementary Material for derivation)

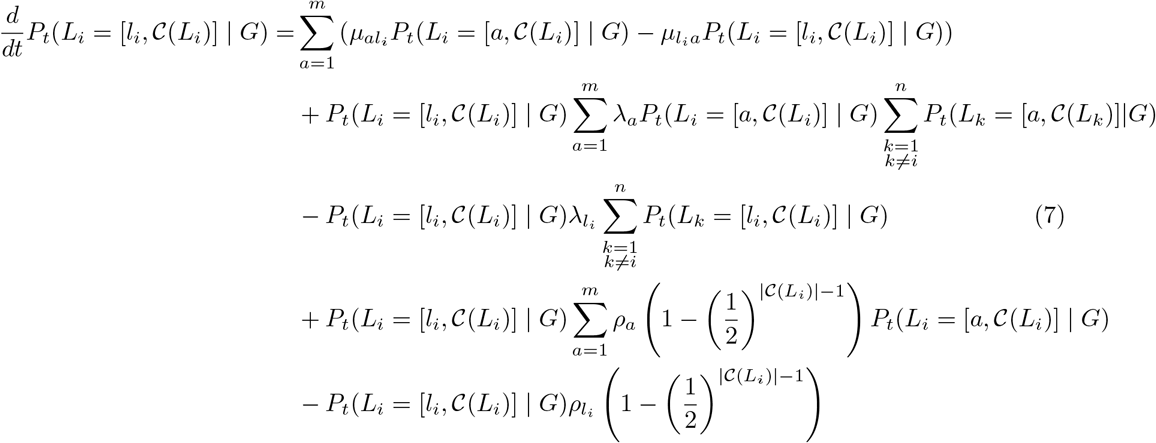

Here, migration, coalescent and reassortment rates (*µ, λ, ρ*) are as described above.

Similarly, we add the necessary reassortment term to the previously derived (Müller, Rasmussen, and Stadler 2018) differential equation for the probability for network history *G* up to time *t* (see Supplementary Material for derivation)

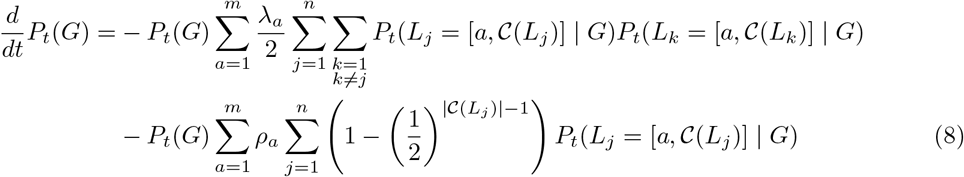

Finally, the above probability will be modified at either coalescent or reassortment events. For the coalescent event between any two lineages *i* and *j*, we have (Müller, Rasmussen, and Stadler 2018)

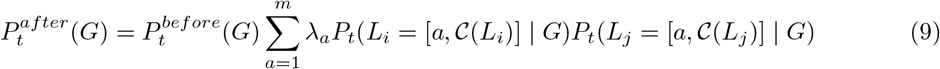

For the reassortment event at a lineage *i* we have

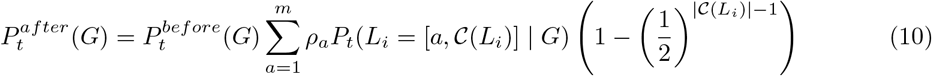

### Numerical integration

To integrate above differential equation numerically, we employ second order Taylor approximation with the numerical integration step size estimated by the third derivative. More details can be found in (Müller, Rasmussen, and Stadler 2018) and the necessary derivatives are given in Supplementary Material.

### Sampling lineage types

We use a stochastic mapping technique (Nielsen 2002; Huelsenbeck, Nielsen, and Bollback 2003) to sample the migration (type change) histories over the network. That is, we first perform backward in time integration, as described above, and obtain the root node type probability distribution ***π*** = (*π*_1_, …, *π*_*m*_). Then, we sample a type *π*_*mrca*_ for the root node from ***π*** and simulate migration as continuous time-inhomogeneous Markov process along the network branches forward in time. The times and types of the endpoints (network leaves) are known. We add an additional constraint, that both parents of a reassortment event must have the same type immediately before joining. If either of the two last conditions are not met, simulated history is rejected. Let *Q* be a time-dependent generator matrix with off-diagonal entries *q*_*ab*_ *>* 0 and diagonal entries 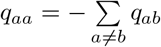. Briefly, the forward simulation algorithm on a network interval between time *t*_0_ and time *t*_1_ is:

- For each coexisting lineage *L*_*i*_ with current type *l*_*i*_ draw an exponential waiting time *τ*_*i*_ ∼ Exponential(*Q*^*i*^).
- If 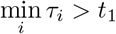, all lineages retain their types. If the node at time *t*_1_ is a reassortment node, we check that its parent lineages have the same simulated type at time *t*_1_. If not, we reject the simulation and start from the root node again. Otherwise, continue with time *t*_0_ := *t*_1_ and *t*_1_ reset as the end of the next network interval.
- If 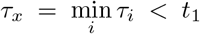 simulate the new type *y* for a lineage *L*_*x*_, from a distribution with probabilities 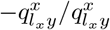, such that *L*_*x*_ configuration changes from [*l*_*x*_, 𝒞(*L*_*x*_)] to [*y*, 𝒞(*L*_*x*_)]. Set *t*_0_ = *τ*_*x*_ and repeat from the first step.

The equations, needed to obtain the generator matrix are given in the Supplementary Material. We implemented this stochastic mapping for both BEAST2 packages, SCoRe and MASCOT.

### Root state conditioning for the structured segment trees and networks

First we will discuss the filtering of the posterior distribution for segment trees, obtained by MAS-COT. Let {*a, g*} be the two possible types of the segment root node and *p*^*i*^_*a*_ - a probability of segment tree *i* having the root node in type *a*. In order to filter the posterior tree distribution, for each pair of segment trees we sample a combination of root types (*aa, ga, ag, gg*) given by the probabilities (*p*^*i*^_*a*_*p*^*j*^_*a*_, *p*^*i*^_*g*_*p*^*j*^_*a*_, *p*^*i*^_*a*_*p*^*j*^_*g*_, *p*^*i*^_*g*_*p*^*j*^_*g*_) and accept the trees if sample is equal to *aa*. Note that the required root type probabilities are already calculated by MASCOT.

In order to filter a posterior distribution of structured networks, obtained by SCoRe, we first obtain the network nodes, corresponding to the root of each segment tree. Then, we accept such network if both nodes were mapped to desired type by stochastic mapping algorithm (see Sampling lineage types above).

### Implementation

SCoRe is a BEAST 2.5 (R. Bouckaert et al. 2018) package for structured coalescent with reassortment network inference and depends on the two packages CoalRe (Müller, Stolz, et al. 2020) and MASCOT (Müller, Rasmussen, and Stadler 2018). Most MCMC operators of CoalRe for an unstructured network proposal can also be used in our case. However, we have to adjust parameter values supplied to the operator which re-simulates unstructured network above the most recent common ancestor of all segment trees as this section of the network is not informed by the sequencing data (see “Gibbs operator” in (Müller, Stolz, et al. 2020) for more details). In the unstructured setting, we may re-simulate with the most recent update of the parameter values. The network proposed by this operator would always be accepted as its Hastings ratio is the inverse ratio for the density of the current and proposed networks.

The source code can be found here: https://github.com/jugne/SCORE and includes tools to obtain summarized MCC networks and investigate reassortment and migration correlations. The MCC networks can be visualised with https://icytree.org (Vaughan 2017) or using an extension of the baltic package (https://github.com/evogytis/baltic): https://github.com/jugne/score-paper-material.git. The tutorial on the installation and usage of the package can be found here: https://github.com/jugne/SCoRe-tutorial.

### Simulation study setup

We have run two well-calibrated simulation studies: (1) assuming known fixed structured coalescent network and inferring its respective effective population sizes, migration and reassortment rates and (2) inferring both, the network and its parameters. Genetic sequences for each segment were simulated according to JC69 (Jukes and Cantor 1969) substitution model. In the fixed network setting, we simulated 1000 networks with 100 taxa, each carrying four segments and being in one out of 2 types. For every simulation, effective population sizes, reassortment and migration rates are randomly drawn from log-normal distributions, 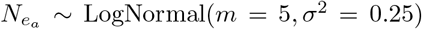, *ρ*_*a*_ ∼ LogNormal(*m* = 0.1, *σ* ^2^ = 0.25), *µ*_*ab*_ ∼ LogNormal(*m* = 0.2, *σ* ^2^ = 0.25), for any type *a, b ∈* {1, 2} and *a* = *b*. Note that here *m* denotes the mean of real variable. The mean for the natural logarithm of this variable can be calculated as 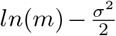. Then, we inferred the parameter values, given the simulated networks, using the above parameter distributions as priors.

For joint network and parameters inference, we simulated 100 networks and embedding of the segment trees for 100 taxa with four segments. The types and sampling times were drawn as described above. clock rates were set to either high (5 × 10^−3^), low (5 × 10^−4^) or mixed (two segments with high and two with low). The prior distribution of a reassortment rate was the same as above, while we set a mean of 2 for the log-normal distribution of effective population sizes. For migration rates, we studied cases where the prior distributions were the same as detailed above and where it was the exponential distribution with mean 0.2. The different migration prior was studied because we noticed it to be a more natural choice when applying the model to the real seasonal influenza dataset. Each inference was run for 48 hours, and we used only those runs for which the effective sample size (ESS) of posterior probability was higher than 100.

Finally, we investigated the relative error in parameter estimates obtained by SCoRe and MASCOT. We again simulated 100 network with 100 taxa and four segments, each simulation repeated with high and low clock rates. The parameters were drawn from the followingdistributions: 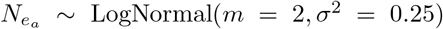, *ρ*_*a*_ ∼ LogNormal(*m* = 0.5, *σ* ^2^ = 0.25), *µ*_*ab*_ ∼ LogNormal(*m* = 0.2, *σ* ^2^ = 0.25), for any type *a, b ∈* {1, 2} and *a* = *b*. Higher reassortment rates were chosen to model a case where reassortment highly influences the evolution of a virus. Both methods were using the same distributions as parameter priors and additionally we set reassortment rate to the true value in SCoRe. The relative error was calculated as 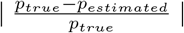, where *p*_*true*_ is the true parameter value and *p*_*estimated*_ is the median parameter estimate obtained by SCoRe or MASCOT.

### Datasets and data availability

We gathered a dataset of 1410 pairs for avian influenza A/H5N1 segment HA and NA sequences for Galliformes (822) and Anseriformes (588) with complete sampling dates between 2008 and 2016 from GISAID (Shu and McCauley 2017, https://www.gisaid.org/) and the Influenza Research Database (Zhang et al. 2017, http://www.fludb.org). Then we iterate over all sampling times and pool sequences that are (1) from the same geographic location, (2) of the same bird order and are within 30 days distance. There we 123 such location-order-date pools for Anseriformes and 159 for Galliformes. Then we randomly sample one sequence per pool and discard the remaining sequences in this pool. Finally, we further reduce this subsample to 100 segment sequence pairs for Anseriformes and Galliformes. The last step is done by weighted subsampling with regard to date and location. We repeat this procedure 10 times, thus obtaining 10 random subsamples of our data. Given random sampling from the location-order-date pools and further random reduction, 10 subsampled datasets may overlap, but are not identical. The BEAST2 XML files which include sequence names can be found on https://github.com/jugne/score-paper-material.git.

## Supporting information

Sequence aquisition IDs

## Acknowledgement

We thank Claire Guinat and Sophie Seidel for valuable comments and discussions. We also thank the authors, originating and submitting laboratories who generously contributed influenza A/H5N1 sequence data to GISAID’s EpiFlu Database (Shu and McCauley 2017) and the Influenza Research Database (Zhang et al. 2017).

NFM is funded by the Swiss National Science Foundation (P2EZP3 191891). US, TGV and TS thank ETH Zürich for funding.

