## Supplementary material for "Joint inference of migration and reassortment patterns for viruses with segment genomes"

### Supplementary Information

May 15, 2021

#### Structured coalescent with reassortment

In this section we derive the exact and approximate equations need to calculate the probability of the structured coalescent with reassortment network. We seek to maintain notation of Müller, D. A. Rasmussen, and Stadler 2017 and Müller, D. Rasmussen, and Stadler 2018 and will not repeat their derivations if the transition is trivial (i.e., exchanging notation for tree to a network).

As in the main text, let $\mathcal{K} := \{L_i = [l_i, \mathcal{C}(L_i)] \mid i \in \{1, 2, ..., n\}\}$ be a specific network $G$ configuration at some time point $t$, where $n$ is the number of coexisting lineages at this time, and $l_i \in \{1, .., m\}$ is the type of lineage $L_i$, taken from a set of $m$ possible network lineage types (e.g., geographic location or virus host type) and $\mathcal{C}(L_i)$ is the set of ancestral segments carried by lineage $L_i$ with cardinality $|\mathcal{C}(L_i)|$. The time $t$ increases backwards until the most recent ancestor (MRCA) of the network, with $t = 0$ being the time of most recent sample. In order to obtain the probability of a network we need to evaluate the joint probability $P_t := P_t(\mathcal{K}, G)$ of configuration $\mathcal{K}$ at time $t$ and network history $G$ until this time at $t = t_{mrca}$. That is, we need to derive (1) the change in this probability over time, within network intervals between any of the four possible network events and (2) the change, caused by coalescent or reassortment events (we condition on sampling events and marginalise over the all possible migration histories).

##### Interval contribution

As in the main text, let $\lambda_a$ be a coalescent rate of type $a$, $\mu_{ab}$ per lineage backwards migration rate from type $a$ to $b$ ($\mu_{aa} = 0$) and $\rho_a$ per lineage reassortment rate of type $a$. If $k_a(\mathcal{K})$ is the number of lineages in type $a$ for some configuration $\mathcal{K}$ then the total coalescent and migration rates are (Müller, D. A. Rasmussen, and Stadler 2017)

$$\mathcal{C} = \sum_{a=1}^{m} \lambda_a \binom{k_a(\mathcal{K})}{2}, \quad \mathcal{M} = \sum_{i=1}^{n} \sum_{a=1}^{m} \mu_{l_i a}. \tag{1}$$

Let an indicator probability $P_t(L_i = [a, \mathcal{C}(L_i)] \mid \mathcal{K}, G) = 1$ if lineage $L_i$ is in type $a$, given configuration $\mathcal{K}$. Then total reassortment rate is

$$\mathcal{R} = \sum_{a=1}^{m} \rho_a \sum_{i=1}^{n} P_t(L_i = [a, \mathcal{C}(L_i)] \mid \mathcal{K}, G) \left(1 - \left(\frac{1}{2}\right)^{|\mathcal{C}(L_i)|-1}\right). \tag{2}$$

As in the main text, $\left(1 - \left(\frac{1}{2}\right)^{\mathcal{C}(L_j)-1}\right)$ is the probability that reassortment event is observable (see also Müller, Stolz, et al. 2019).

Further, denote $\mathcal{K}_{l_i=a} := \{L_i = [a, \mathcal{C}(L_i)] \cup \{L_j = [l_j, \mathcal{C}(L_j)] \mid j \in \{1, 2, ..., n\}, j \neq i, l_j \in \{1, 2, ..., m\}\}\}$ a network configuration, where lineage $L_i$ is in a type $a$. To obtain the change in $P_t$ over time, we need to evaluate $P_{t+\Delta t}$ based on $P_t$

$$\begin{aligned} P_{t+\Delta t}(\mathcal{K}, G) =& P_t(\mathcal{K}, G)(1 - \mathcal{M}\Delta t - \mathcal{C}\Delta t - \mathcal{R}\Delta t) \\ &+ \sum_{i=1}^{n} \sum_{a=1}^{m} (\mu_{a l_i} \Delta t P_t(\mathcal{K}_{l_i=a}, G)) + O((\Delta t)^2) \end{aligned}$$

The multiplication of the first two terms in the right hand side accounts for no migration, coalescent or reassortment events within the time $\Delta t$. The double summation accounts for any lineage migrating into any type. Subtracting $P_t(\mathcal{K}, G)$ from both sides of the equation, dividing them by $\Delta t$ and letting $\Delta t \longrightarrow 0$ yields

$$\frac{\mathrm{d}P_t(\mathcal{K}, G)}{\mathrm{d}t} = -(\mathcal{M} + \mathcal{C} + \mathcal{R})P_t(\mathcal{K}, G) + \sum_{i=1}^{n} \sum_{a=1}^{m} (\mu_{a l_i} P_t(\mathcal{K}_{l_i=a}, G)).$$

Finally, substituting expressions for $\mathcal{M}, \mathcal{C}, \mathcal{R}$ from equations (1), (2) constitutes an exact differential equation for the change in network configuration probability over time:

$$\begin{aligned}\frac{\mathrm{d}P_t(\mathcal{K},G)}{\mathrm{d}t} =& \sum_{i=1}^{n}\sum_{a=1}^{m}(\mu_{al_i}P_t(\mathcal{K}_{l_i=a},G) - \mu_{al_i}P_t(\mathcal{K},G)) \\ &- \sum_{a=1}^{m}\lambda_a \binom{k_a(\mathcal{K})}{2} P_t(\mathcal{K},G) \qquad (3) \\ &- \sum_{a=1}^{m}\rho_a \sum_{i=1}^{n} P_t(L_i = [a, \mathcal{C}(L_i)] \mid \mathcal{K}, G) \left(1 - \left(\frac{1}{2}\right)^{|\mathcal{C}(L_i)|-1}\right) P_t(\mathcal{K},G) \qquad (4)\end{aligned}$$

As discussed in the main text, there are $m^n$ possible configurations $\mathcal{K}$. It is computationally unfeasible to evaluate all of them, if there are more than a few lineage or types.

Next, we apply the MASCO approximation to the reassortment part of equation (3) (derivation for coalescent and migration parts is equivalent to the tree case, given by Müller, D. A. Rasmussen, and Stadler 2017). Let $\mathcal{K}_{\setminus i} := \{L_j = [l_j, \mathcal{C}(L_j)] \mid j \in \{1,2,...,n\}, j \neq i, l_j \in \{1,2,...,m\}\}$ be a configuration for all lineages, except lineage $i$. Then we can write the marginal lineage type probability

$$P_t(L_i = [l_i, \mathcal{C}(L_i)], G) = \sum_{\mathcal{K}_{\setminus i}} P_t(\mathcal{K},G) \qquad (5)$$

Applying the above summation to the reassortment part of equation (3) yields

$$\sum_{\mathcal{K}_{\setminus i}} \mathcal{R} P_t(\mathcal{K},G) = \sum_{\mathcal{K}_{\setminus i}} \sum_{a=1}^{m}\rho_a \sum_{j=1}^{n} P_t(L_j = [a, \mathcal{C}(L_j)] \mid \mathcal{K}, G) \left(1 - \left(\frac{1}{2}\right)^{|\mathcal{C}(L_j)|-1}\right) P_t(\mathcal{K},G)$$

Separating the summation involving lineage $i$ we get

$$\begin{aligned}\sum_{\mathcal{K}_{\setminus i}} \mathcal{R} P_t(\mathcal{K},G) =& \sum_{a=1}^{m}\rho_a \sum_{\substack{j=1 \\ j\neq i}}^{n} \left(1 - \left(\frac{1}{2}\right)^{|\mathcal{C}(L_i)|-1}\right) \sum_{\mathcal{K}_{\setminus i}} P_t(L_j = [a, \mathcal{C}(L_j)] \mid \mathcal{K}, G) P_t(\mathcal{K},G) \\ &+ \sum_{a=1}^{m}\rho_a \left(1 - \left(\frac{1}{2}\right)^{|\mathcal{C}(L_i)|-1}\right) \sum_{\mathcal{K}_{\setminus i}} P_t(L_i = [a, \mathcal{C}(L_i)] \mid \mathcal{K}, G) P_t(\mathcal{K},G) \qquad (6)\end{aligned}$$

Using equation (5) and allowing for conditional lineage type independence, we can write

$$\begin{aligned}\sum_{\mathcal{K}_{\setminus i}} P_t(L_j = a \mid \mathcal{K}, G) P_t(\mathcal{K},G) &= \sum_{\mathcal{K}_{\setminus i}} P_t(L_j = a, \mathcal{K}, G) \\ &= P_t(L_j = [a, \mathcal{C}(L_j)], L_i = [l_i, \mathcal{C}(L_i)], G) \\ &= P_t(L_j = [a, \mathcal{C}(L_j)], L_i = [l_i, \mathcal{C}(L_i)]) P_t(G) \\ &\overset{MASCO}{=} P_t(L_j = [a, \mathcal{C}(L_j)]) P_t(L_i = [l_i, \mathcal{C}(L_i)], G).\end{aligned}$$

Now, we can use the above to approximate equation (6):

$$\begin{aligned}\sum_{\mathcal{K}_{\setminus i}} \mathcal{R} P_t(\mathcal{K},G) =& \sum_{a=1}^{m}\rho_a \sum_{\substack{j=1 \\ j\neq i}}^{n} \left(1 - \left(\frac{1}{2}\right)^{|\mathcal{C}(L_j)|-1}\right) P_t(L_j = [a, \mathcal{C}(L_j)], L_i = [l_i, \mathcal{C}(L_i)] \mid G) P_t(G) \\ &+ \sum_{a=1}^{m}\rho_a \left(1 - \left(\frac{1}{2}\right)^{|\mathcal{C}(L_i)|-1}\right) P_t(L_i = [a, \mathcal{C}(L_i)], L_i = [l_i, \mathcal{C}(L_i)] \mid G) P_t(G) \\ \overset{MASCO}{=}& P_t(L_i = [l_i, \mathcal{C}(L_i)]\, G) \sum_{a=1}^{m}\rho_a \sum_{\substack{j=1 \\ j\neq i}}^{n} \left(1 - \left(\frac{1}{2}\right)^{|\mathcal{C}(L_j)|-1}\right) P_t(L_j = [a, \mathcal{C}(L_j)] \mid G) \\ &+ P_t(L_i = [l_i, \mathcal{C}(L_i)], G) \rho_{l_i} \left(1 - \left(\frac{1}{2}\right)^{|\mathcal{C}(L_i)|-1}\right) \qquad (7)\end{aligned}$$

Finally, we subtract the above equation (7) from the right hand side of equation (6) in the Supplemental Material of (Müller, D. A. Rasmussen, and Stadler 2017) and obtain the full ODE for the change in marginal lineage type probability over time:

$$
\begin{aligned}
\frac{d}{dt}P_t(L_i=[l_i,\mathcal{C}(L_i)],G) =& \sum_{a=1}^{m}\left(\mu_{al_i}P_t(L_i=[a,\mathcal{C}(L_i)],) - \mu_{l_ia}P_t(L_i=[l_i,\mathcal{C}(L_i)],G)\right)\\
&- P_t(L_i=[l_i,\mathcal{C}(L_i)],G)\sum_{a=1}^{m}\frac{\lambda_a}{2}\sum_{\substack{j=1\\j\neq i}}^{n}\sum_{\substack{k=1\\k\neq j,i}}^{n}P_t(L_j=[a,\mathcal{C}(L_j)]\mid G)P_t(L_k=[a,\mathcal{C}(L_k)]\mid G)\\
&- P_t(L_i=[l_i,\mathcal{C}(L_i)],G)\lambda_{l_i}\sum_{\substack{k=1\\k\neq i}}^{n}P_t(L_k=[l_i,\mathcal{C}(L_k)\mid G)\\
&- P_t(L_i=[l_i,\mathcal{C}(L_i)],G)\sum_{a=1}^{m}\rho_a\sum_{\substack{j=1\\j\neq i}}^{n}\left(1-\left(\frac{1}{2}\right)^{|\mathcal{C}(L_j)|-1}\right)P_t(L_j=[a,\mathcal{C}(L_j)]\mid G)\\
&- P_t(L_i=[l_i,\mathcal{C}(L_i)],G)\rho_{l_i}\left(1-\left(\frac{1}{2}\right)^{|\mathcal{C}(L_i)|-1}\right) \qquad (8)
\end{aligned}
$$

**Reassortment event contribution**

The exact and approximate forms of the coalescent event contribution are equivalent to the tree case (Müller, D. A. Rasmussen, and Stadler 2017), with the appropriate segment union of child lineages, explained in the main text. After the reassortment event at lineage $i$ of type $a$, configuration $\mathcal{K}$ changes to $\mathcal{K}' := \{\{L_{p1}=[a,\mathcal{C}(L_{p1})], L_{p2}=[a,\mathcal{C}(L_{p2})] \mid p1,p2\in\{1,...,n+1\}\}\cup\{L_s=[l_s,\mathcal{C}(L_s)] \mid s\in\{1,...,n+1\}, s\notin\{p1,p2\}, l_s\in\{1,...,m\}\}$, and $\mathcal{C}(L_{p1})\cup\mathcal{C}(L_{p1})=\mathcal{C}(L_i)$, $\mathcal{C}(L_{p1})\cap\mathcal{C}(L_{p1})=\emptyset$. Then, in the exact form, such reassortment event changes the probability $P_t$ as follows:

$$
P_t^{after}(G) = P_t^{before}(G)P_t(\mathcal{K}_{l_i=a}\mid G)\rho_a\left(1-\left(\frac{1}{2}\right)^{|\mathcal{C}(L_i)|-1}\right)
$$

Applying the MASCO approximation we get the equation (10) in the main text.

### Numerical Integration

As discused by Müller, D. Rasmussen, and Stadler 2018 evaluating derivative (8) poses computational difficulties due to ever decreasing values. The authors there suggest to overcome this by evaluating conditional probability $\frac{d}{dt}P_t(L_i=[l_i,\mathcal{C}(L_i)]\mid G)$. They noted that it can be taking the following difference:

$$
\frac{d}{dt}P_t(L_i=[l_i,\mathcal{C}(L_i)],G) = \frac{d}{dt}\frac{P_t(L_i=[l_i,\mathcal{C}(L_i)],G)}{P_t(G)} = \frac{\frac{dt}{dt}P_t(L_i=[l_i,\mathcal{C}(L_i)],G)}{P_t(G)} - \frac{P_t(L_i=[l_i,\mathcal{C}(L_i)]\mid G)\frac{d}{dt}P_t(G)}{P_t(G)}
$$

We use the same method to obtain the expression for $\frac{d}{dt}P_t(L_i=[l_i,\mathcal{C}(L_i)]\mid G)$ when reassortment is included. First, we divide the equation (8) by $P_t(G)$

$$
\begin{aligned}
\frac{\frac{d}{dt}P_t(L_i=[l_i,\mathcal{C}(L_i)],G)}{P_t(G)} =& \sum_{a=1}^{m}\left(\mu_{al_i}P_t(L_i=[a,\mathcal{C}(L_i)]\mid G) - \mu_{l_ia}P_t(L_i=[l_i,\mathcal{C}(L_i)]\mid G)\right)\\
&- P_t(L_i=[l_i,\mathcal{C}(L_i)]\mid G)\sum_{a=1}^{m}\frac{\lambda_a}{2}\sum_{\substack{j=1\\j\neq i}}^{n}\sum_{\substack{k=1\\k\neq j,i}}^{n}P_t(L_j=[a,\mathcal{C}(L_j)]\mid G)P_t(L_k=[a,\mathcal{C}(L_k)]\mid G)\\
&- P_t(L_i=[l_i,\mathcal{C}(L_i)]\mid G)\lambda_{l_i}\sum_{\substack{k=1\\k\neq i}}^{n}P_t(L_k=[l_i,\mathcal{C}(L_k)\mid G)\\
&- P_t(L_i=[l_i,\mathcal{C}(L_i)]\mid G)\sum_{a=1}^{m}\rho_a\sum_{\substack{j=1\\j\neq i}}^{n}\left(1-\left(\frac{1}{2}\right)^{|\mathcal{C}(L_j)|-1}\right)P_t(L_j=[a,\mathcal{C}(L_j)]\mid G)\\
&- P_t(L_i=[l_i,\mathcal{C}(L_i)]\mid G)\rho_{l_i}\left(1-\left(\frac{1}{2}\right)^{|\mathcal{C}(L_i)|-1}\right) \qquad (9)
\end{aligned}
$$

The second term of the difference above is

$$\frac{P_t(L_i = [l_i, \mathcal{C}(L_i)] \mid G)\frac{d}{dt}P_t(G)}{P_t(G)} = -\frac{P_t(L_i = [l_i, \mathcal{C}(L_i)] \mid G)}{P_t(G)}\Bigg[P_t(G)\sum_{b=1}^{m}\frac{\lambda_b}{2}\sum_{\substack{j=1\\j\neq i}}^{n}\sum_{\substack{k=1\\k\neq j,i}}^{n}P_t(L_j = [b, \mathcal{C}(L_j)] \mid G)P_t(L_k = b|G)$$
$$+ P_t(G)\sum_{a=1}^{m}P_t(L_i = [a, \mathcal{C}(L_i)] \mid G)\lambda_a\sum_{\substack{k=1\\k\neq i}}^{n}P_t(L_k = [a, \mathcal{C}(L_k)] \mid G)$$
$$+ P_t(G)\sum_{b=1}^{m}\rho_b\sum_{\substack{j=1\\j\neq i}}^{n}\left(1 - \left(\frac{1}{2}\right)^{|\mathcal{C}(L_j)|-1}\right)P_t(L_j = [b, \mathcal{C}(L_j)] \mid G)$$
$$+ P_t(G)\sum_{a=1}^{m}P_t(L_i = [a, \mathcal{C}(L_i)] \mid G)\rho_a\left(1 - \left(\frac{1}{2}\right)^{|\mathcal{C}(L_i)|-1}\right)\Bigg]$$
$$= -P_t(L_i = [l_i, \mathcal{C}(L_i)] \mid G)\sum_{a=1}^{m}\frac{\lambda_a}{2}\sum_{j=1}^{n}\sum_{\substack{k=1\\k\neq j}}P_t(L_j = [a, \mathcal{C}(L_j)] \mid G)P_t(L_k = [a, \mathcal{C}(L_k)] \mid G)$$
$$- P_t(L_i = [l_i, \mathcal{C}(L_i)] \mid G)\sum_{a=1}^{m}\rho_a\sum_{j=1}^{n}\left(1 - \left(\frac{1}{2}\right)^{|\mathcal{C}(L_j)|-1}\right)P_t(L_j = [a, \mathcal{C}(L_j)] \mid G) \tag{10}$$

Subtracting (10) from (9) yields the derivative (7) in the main text

$$\frac{d}{dt}P_t(L_i = [l_i, \mathcal{C}(L_i)] \mid G) = \sum_{a=1}^{m}\left(\mu_{al_i}P_t(L_i = [a, \mathcal{C}(L_i)] \mid G) - \mu_{l_ia}P_t(L_i = [l_i, \mathcal{C}(L_i)] \mid G)\right)$$
$$+ P_t(L_i = [l_i, \mathcal{C}(L_i)] \mid G)\sum_{a=1}^{m}\lambda_a P_t(L_i = [a, \mathcal{C}(L_i)] \mid G)\sum_{\substack{k=1\\k\neq i}}^{n}P_t(L_k = [a, \mathcal{C}(L_k)]|G)$$
$$- P_t(L_i = [l_i, \mathcal{C}(L_i)] \mid G)\lambda_{l_i}\sum_{\substack{k=1\\k\neq i}}^{n}P_t(L_k = [l_i, \mathcal{C}(L_i)] \mid G) \tag{11}$$
$$+ P_t(L_i = [l_i, \mathcal{C}(L_i)] \mid G)\sum_{a=1}^{m}\rho_a\left(1 - \left(\frac{1}{2}\right)^{|\mathcal{C}(L_i)|-1}\right)P_t(L_i = [a, \mathcal{C}(L_i)] \mid G)$$
$$- P_t(L_i = [l_i, \mathcal{C}(L_i)] \mid G)\rho_{l_i}\left(1 - \left(\frac{1}{2}\right)^{|\mathcal{C}(L_i)|-1}\right)$$

The derivative of probability $P_t(G)$ for a network $G$ up to time $t$ is obtained from the derivative of joint lineage type and network probability. We know that

$$\frac{d}{dt}P_t(G) = \sum_{a=1}^{m}\frac{d}{dt}P_t(L_i = [a, \mathcal{C}(L_i)], G) = \frac{d}{dt}\sum_{a=1}^{m}P_t(L_i = [a, \mathcal{C}(L_i)], G)$$

To get the right hand side of the above expression we multiply equation (10) by $\frac{P_t(G)}{P_t(L_i = l_i|G)}$:

$$\frac{d}{dt}P_t(G) = -P_t(G)\sum_{a=1}^{m}\frac{\lambda_a}{2}\sum_{j=1}^{n}\sum_{\substack{k=1\\k\neq j}}P_t(L_j = [a, \mathcal{C}(L_j)] \mid G)P_t(L_k = a|G)$$
$$- P_t(G)\sum_{a=1}^{m}\rho_a\sum_{j=1}^{n}\left(1 - \left(\frac{1}{2}\right)^{|\mathcal{C}(L_j)|-1}\right)P_t(L_j = [a, \mathcal{C}(L_j)] \mid G)$$

##### Taylor series approximation

We use Taylor series approximation of the second order with the third derivative used to estimate step size (see Müller, D. Rasmussen, and Stadler 2018 for more details). Therefore, we need an expression for the exact second derivative of $P_t(L_i = [l_i, \mathcal{C}(L_i)] \mid G)$ and approximate expression for its third derivative.

Second derivative is obtained by applying the product rule to equation (11):

$$
\begin{aligned}
\frac{d^2 P_t(L_i = [l_i, \mathcal{C}(L_i)] \mid G)}{dt^2} =& \sum_{a=1}^{m} \left( \mu_{al_i} \frac{d}{dt} P_t(L_i = [a, \mathcal{C}(L_i)] \mid G) - \mu_{l_i a} \frac{d}{dt} P_t(L_i = [l_i, \mathcal{C}(L_i)] \mid G) \right) \\
&+ \frac{d}{dt} P_t(L_i = [l_i, \mathcal{C}(L_i)] \mid G) \sum_{a=1}^{m} \lambda_a P_t(L_i = [a, \mathcal{C}(L_i)] \mid G) \sum_{\substack{k=1 \\ k \neq i}}^{n} P_t(L_k = [a, \mathcal{C}(L_k)] \mid G) \\
&+ P_t(L_i = [l_i, \mathcal{C}(L_i)] \mid G) \sum_{a=1}^{m} \lambda_a \frac{d}{dt} P_t(L_i = [a, \mathcal{C}(L_i)] \mid G) \sum_{\substack{k=1 \\ k \neq i}}^{n} P_t(L_k = [a, \mathcal{C}(L_k)] \mid G) \\
&+ P_t(L_i = [l_i, \mathcal{C}(L_i)] \mid G) \sum_{a=1}^{m} \lambda_a P_t(L_i = [a, \mathcal{C}(L_i)] \mid G) \sum_{\substack{k=1 \\ k \neq i}}^{n} \frac{d}{dt} P_t(L_k = [a, \mathcal{C}(L_k)] \mid G) \\
&- \frac{d}{dt} P_t(L_i = [l_i, \mathcal{C}(L_i)] \mid G) \lambda_{l_i} \sum_{\substack{j=1 \\ j \neq i}}^{n} P_t(L_k = [l_i, \mathcal{C}(L_k)] \mid G) \\
&- P_t(L_i = [l_i, \mathcal{C}(L_i)] \mid G) \lambda_{l_i} \sum_{\substack{j=1 \\ j \neq i}}^{n} \frac{d}{dt} P_t(L_k = [l_i, \mathcal{C}(L_k)] \mid G) \\
&+ \frac{d}{dt} P_t(L_i = [l_i, \mathcal{C}(L_i)] \mid G) \sum_{a=1}^{m} \rho_a \left( 1 - \left( \frac{1}{2} \right)^{|\mathcal{C}(L_i)|-1} \right) P_t(L_i = [a, \mathcal{C}(L_i)] \mid G) \\
&+ P_t(L_i = [l_i, \mathcal{C}(L_i)] \mid G) \sum_{a=1}^{m} \rho_a \left( 1 - \left( \frac{1}{2} \right)^{|\mathcal{C}(L_i)|-1} \right) \frac{d}{dt} P_t(L_i = [a, \mathcal{C}(L_i)] \mid G) \\
&- \frac{d}{dt} P_t(L_i = [l_i, \mathcal{C}(L_i)] \mid G) \rho_{l_i} \left( 1 - \left( \frac{1}{2} \right)^{|\mathcal{C}(L_i)|-1} \right)
\end{aligned} \tag{12}
$$

The third derivative is used to determine the step size $\Delta t$ of integration and we evaluate it approximately. We use the same assumptions as those, made by Müller, D. Rasmussen, and Stadler 2018:

- the sum of probability mass in a type over all lineages but lineage $i$ does not change;
- the sum of the derivatives of lineage $i$ coalescing in any type does not change.

Then the third and fourth lines of the second derivative (12) are equal to zero and

$$
\begin{aligned}
\frac{d^3 P_t(L_i = [l_i, \mathcal{C}(L_i)] \mid G)}{dt^3} \approx& \sum_{a=1}^{m} \left( \mu_{al_i} \frac{d^2}{dt^2} P_t(L_i = [a, \mathcal{C}(L_i)] \mid G) - \mu_{l_i a} \frac{d^2}{dt^2} P_t(L_i = [l_i, \mathcal{C}(L_i)] \mid G) \right) \\
&+ \frac{d^2}{dt^2} P_t(L_i = [l_i, \mathcal{C}(L_i)] \mid G) \left( \sum_{a=1}^{m} \lambda_a P_t(L_i = [a, \mathcal{C}(L_i)] \mid G) \sum_{\substack{k=1 \\ k \neq i}}^{n} P_t(L_k = [a, \mathcal{C}(L_k)] \mid G) \right) \\
&- \frac{d^2}{dt^2} P_t(L_i = [l_i, \mathcal{C}(L_i)] \mid G) \lambda_{l_i} \sum_{\substack{k=1 \\ k \neq i}}^{n} P_t(L_k = [l_i, \mathcal{C}(L_k)] \mid G) \\
&+ \frac{d^2}{dt^2} P_t(L_i = [l_i, \mathcal{C}(L_i)] \mid G) \left( \sum_{a=1}^{m} \rho_a \left( 1 - \left( \frac{1}{2} \right)^{|\mathcal{C}(L_i)|-1} \right) P_t(L_i = [a, \mathcal{C}(L_i)] \mid G) \right) \\
&+ 2 \frac{d}{dt} P_t(L_i = [l_i, \mathcal{C}(L_i)] \mid G) \left( \sum_{a=1}^{m} \rho_a \left( 1 - \left( \frac{1}{2} \right)^{|\mathcal{C}(L_i)|-1} \right) \frac{d}{dt} P_t(L_i = [a, \mathcal{C}(L_i)] \mid G) \right) \\
&+ P_t(L_i = [l_i, \mathcal{C}(L_i)] \mid G) \left( \sum_{a=1}^{m} \rho_a \left( 1 - \left( \frac{1}{2} \right)^{|\mathcal{C}(L_i)|-1} \right) \frac{d^2}{dt^2} P_t(L_i = [a, \mathcal{C}(L_i)] \mid G) \right) \\
&- \frac{d^2}{dt^2} P_t(L_i = [l_i, \mathcal{C}(L_i)] \mid G) \rho_{l_i} \left( 1 - \left( \frac{1}{2} \right)^{|\mathcal{C}(L_i)|-1} \right)
\end{aligned} \tag{13}
$$

#### Stochastic mapping of lineage types

As discussed in the main text, we seek to show that lineage type change follows a continuous Markov process and obtain the expression for its generator matrix. Let $\mathcal{K}$ be network lineage type configuration as above at time $t$, $\mathcal{K}_{mrca}$ be the configuration at the network root, and $\mathcal{K}_f$ a configuration of network leaf nodes. Further, define $G_{>t}$ and $G_{<t}$ to be the parts of the network respectively older and younger than $t$. Then, we can write

$$P(\mathcal{K} \mid G, \mathcal{K}^f) = \frac{\sum_{\mathcal{K}_{mrca}} P(\mathcal{K}_{mrca}, G_{<t} \mid \mathcal{K}, G_{>t}) P(\mathcal{K}, G_{>t} \mid \mathcal{K}_f)}{\sum_{\mathcal{K}'_{mrca}} \sum_{\mathcal{K}'} P(\mathcal{K}'_{mrca}, G_{<t} \mid \mathcal{K}', G_{>t}) P(\mathcal{K}, G_{>t} \mid \mathcal{K}'_f)}.$$

For stochastic mapping algorithm, we seek to evaluate the lineage type probability, given the network $G$, its configuration at the time of the root $\mathcal{K}_{mrca}$ and at the time of the leave nodes $\mathcal{K}_f$:

$$P(\mathcal{K} \mid G, \mathcal{K}_{mrca}, \mathcal{K}_f)) = \frac{P(\mathcal{K}_{mrca}, G_{<t} | \mathcal{K}, G_{>t}) P(\mathcal{K}, G_{>t} \mid \mathcal{K}_f)}{\sum_{\mathcal{K}'} P(\mathcal{K}'_{mrca}, G_{<t} | \mathcal{K}', G_{>t}) P(\mathcal{K}', G_{>t} \mid \mathcal{K}'_f)}. \tag{14}$$

In order to derive the stochastic mapping equation we need to derive the Kolmogorov backward equations for $P(\mathcal{K}, G_{>t} \mid \mathcal{K}_f)$. Kolmogorov forward equation is given by (3) above. Corresponding backward equation can be derived by noting that $P(\mathcal{K}, G_{>t} \mid \mathcal{K}_f)$ does not depend on $z$ and we can express the process using intermediate types:

$$\begin{aligned}
\frac{d}{dz} P(\mathcal{K}, G_{>t} \mid \mathcal{K}_f) = 0 &= \frac{d}{dz} \sum_{\mathcal{K}_z} P(\mathcal{K}, G_{<z} | \mathcal{K}_z, G_{>z}) P(\mathcal{K}_z, G_{>z} \mid \mathcal{K}_f) \\
&= \sum_{\mathcal{K}_z} \left[ \frac{d}{dz} P(\mathcal{K}, G_{<z} | \mathcal{K}_z, G_{>z}) \right] P(\mathcal{K}_z, G_{>z} \mid \mathcal{K}_f) \\
&+ \sum_{\mathcal{K}_z} P(\mathcal{K}, G_{<z} | \mathcal{K}_z, G_{>z}) \left[ \frac{d}{dz} P(\mathcal{K}_z, G_{>z} | \mathcal{K}_f) \right] \\
&= S_1 + S_2.
\end{aligned}$$

Substituting the forward equation into $S_2$, we have

$$\begin{aligned}
S_2 = &- \sum_{\mathcal{K}_z} P(\mathcal{K}, G_{<z} | \mathcal{K}_z, G_{>z}) \sum_{i=1}^{n_z} \sum_{a=1}^{m} [\mu_{a l_i} P(\mathcal{K}_{z, l_i = a}, G_{>z} \mid \mathcal{K}_f) - \mu_{l_i a} P(\mathcal{K}_z, G_{>z} | \mathcal{K}_f)] \\
&+ \sum_{\mathcal{K}_z} P(\mathcal{K}, G_{<z} | \mathcal{K}_z, G_{>z}) \sum_{a=1}^{m} \lambda_a \binom{k_a(\mathcal{K}_z)}{2} P(\mathcal{K}_z, G_{>z} | \mathcal{K}_f) \\
&+ \sum_{\mathcal{K}_z} P(\mathcal{K}, G_{<z} | \mathcal{K}_z, G_{>z}) \sum_{a=1}^{m} \rho_a \sum_{j}^{n_n} P(L_j = [a, \mathcal{C}(L_j)] \mid \mathcal{K}, G_{>z}) \left( 1 - \left( \frac{1}{2} \right)^{|\mathcal{C}(L_j)| - 1} \right) P(\mathcal{K}_z, G_{>z} | \mathcal{K}_f) \\
= &- \sum_{\mathcal{K}_z} P(\mathcal{K}_z, G_{>z} | \mathcal{K}_f) \sum_{i=1}^{n_z} \sum_{a=1}^{m} \mu_{l_i a} [P(\mathcal{K}, G_{<z} \mid \mathcal{K}_{z, l_i = a}, G_{>z}) - P(\mathcal{K}, G_{<z} | \mathcal{K}_z, G_{>z})] \\
&+ \sum_{\mathcal{K}_z} P(\mathcal{K}_z, G_{>z} | \mathcal{K}_f) \sum_{a=1}^{m} \lambda_a \binom{k_a(\mathcal{K}_z)}{2} P(\mathcal{K}, G_{<z} | \mathcal{K}_z, G_{>z}) \\
&+ \sum_{\mathcal{K}_z} P(\mathcal{K}_z, G_{>z} | \mathcal{K}_f) \sum_{a=1}^{m} \rho_a \sum_{j}^{n_z} P(L_j = [a, \mathcal{C}(L_j)] \mid \mathcal{K}, G_{>z}) \left( 1 - \left( \frac{1}{2} \right)^{|\mathcal{C}(L_j)| - 1} \right) P(\mathcal{K}, G_{<z} | \mathcal{K}_z, G_{>z}).
\end{aligned}$$

Since $S_1 = -S_2$, we get

$$\begin{aligned}
\frac{d}{dz} P(\mathcal{K}, G_{<z} | \mathcal{K}_z, G_{>z}) = &\sum_{i=1}^{|\mathcal{K}_z|} \sum_{a=1}^{m} \mu_{l_i a} [P(\mathcal{K}, G_{<z} | \mathcal{K}_{z, l_i = a}, G_{>z}) - P(\mathcal{K}, G_{<z} | \mathcal{K}_z, G_{>z})] \\
&- \sum_{a=1}^{m} \lambda_a \binom{k_a(\mathcal{K}_z)}{2} P(\mathcal{K}, G_{<z} | \mathcal{K}_z, G_{>z}) \\
&- \sum_{a=1}^{m} \rho_a \sum_{j}^{|\mathcal{K}_z|} P(L_j = [a, \mathcal{C}(L_j)] \mid \mathcal{K}, G_{>z})(\mathcal{K}) \left( 1 - \left( \frac{1}{2} \right)^{|\mathcal{C}(L_j)| - 1} \right) P(\mathcal{K}, G_{<z} | \mathcal{K}_z, G_{>z}).
\end{aligned}$$

Using the above, we can evaluate equation (14)

$$\frac{d}{dt}P(\mathcal{K}|G,\mathcal{K}_{mrca},\mathcal{K}_f) = \frac{1}{P(\mathcal{K}_{mrca},G|\mathcal{K}_f)}\left[\left(\frac{d}{dt}P(\mathcal{K}_{mrca},G_{<t}|\mathcal{K},G_{>t})\right)P(\mathcal{K},G_{>t}|\mathcal{K}_f)\right]$$

$$+ P(\mathcal{K}_{mrca},G_{<t}|\mathcal{K},G_{>t})\left(\frac{d}{dz}P(\mathcal{K},G_{>t}|\mathcal{K}_f)\right)$$

$$=\frac{1}{P(\mathcal{K}_{mrca},G|\mathcal{K}_f)}[\sum_{i=1}^{n}\sum_{a=1}^{m}\mu_{l_i a}\left[P(\mathcal{K}_{mrca},G_{<t}|\mathcal{K}_{l_i=a},G_{>t}) - P(\mathcal{K}_{mrca},G_{<t}|\mathcal{K},G_{>t})\right]P(\mathcal{K},G_{>t}|\mathcal{K}_f)$$

$$-\sum_{a=1}^{m}\lambda_a\binom{k_a(\mathcal{K})}{2}P(\mathcal{K}_{mrca},G_{<t}|\mathcal{K},G_{>t})P(\mathcal{K},G_{>t}|\mathcal{K}_f)$$

$$-\sum_{a=1}^{m}\rho_a\sum_{j}^{n}P(L_j=[a,\mathcal{C}(L_j)]\mid\mathcal{K},G_{>z})(\mathcal{K})\left(1-\left(\frac{1}{2}\right)^{|\mathcal{C}(L_j)|-1}\right)P(\mathcal{K}_{mrca},G_{<t}|\mathcal{K},G_{>t})P(\mathcal{K},G_{>t}|\mathcal{K}_f)$$

$$-\sum_{i=1}^{n}\sum_{a=1}^{m}P(\mathcal{K}_{mrca},G_{<t}|\mathcal{K},G_{>t})\left[\mu_{al_i}P(\mathcal{K}_{l_i=a},G_{>t}|\mathcal{K}_f - \mu_{l_i a}P(\mathcal{K},G_{>t}|\mathcal{K}_f)\right]$$

$$+\sum_{a=1}^{m}\lambda_a\binom{k_a(\mathcal{K})}{2}P(\mathcal{K}_{mrca},G_{<t}|\mathcal{K},G_{>t})P(\mathcal{K},G_{>t}|\mathcal{K}_f)$$

$$+\sum_{a=1}^{m}\rho_a\sum_{j=1}^{n}P(L_j=[a,\mathcal{C}(L_j)]\mid\mathcal{K},G_{>z})(\mathcal{K})\left(1-\left(\frac{1}{2}\right)^{|\mathcal{C}(L_j)|-1}\right)P(\mathcal{K}_{mrca},G_{<t}|\mathcal{K},G_{>t})P(\mathcal{K},G_{>t}|\mathcal{K}_f)]$$

Cancelling the reassortment and coalescent terms, we obtain

$$\frac{d}{dt}P(\mathcal{K}|G,\mathcal{K}_{mrca},\mathcal{K}_f) = \frac{1}{P(\mathcal{K}_{mrca},G|\mathcal{K}_f)}\Bigg[$$

$$\sum_{i=1}^{n}\sum_{a=1}^{m}\mu_{l_i a}\left[P(\mathcal{K}_{mrca},G_{<t}|\mathcal{K}_{l_i=a},G_{>t}) - P(\mathcal{K}_{mrca},G_{<t}|\mathcal{K},G_{>t})\right]P(\mathcal{K},G_{>t}|\mathcal{K}_f)$$

$$-\sum_{i=1}^{n}\sum_{a=1}^{m}P(\mathcal{K}_{mrca},G_{<t}|\mathcal{K},G_{>t})\left[\mu_{al_i}P(\mathcal{K}_{l_i=a},G_{>t}|\mathcal{K}_f) - \mu_{l_i a}P(\mathcal{K},G_{>t}|\mathcal{K}_f)\right]\Bigg]$$

$$=\frac{1}{P(\mathcal{K}_{mrca},G|\mathcal{K}_f)}\Bigg[\sum_{i=1}^{n}\sum_{a=1}^{m}\Big(\mu_{l_i a}P(\mathcal{K}_{mrca},G_{<t}|\mathcal{K}_{l_i=a},G_{>t})P(\mathcal{K},G_{>t}|\mathcal{K}_f)$$

$$-\mu_{al_i}P(\mathcal{K}_{mrca},G_{<t}|\mathcal{K},G_{>t})P(\mathcal{K}_{l_i=a},G_{>t}|\mathcal{K}_f)\Big)\Bigg]$$

Let $q^i_{ab} = \mu_{ba}\frac{P(\mathcal{K}_{l_i=b},G_{>t}|\mathcal{K}_f)}{P(\mathcal{K}_{l_i=a},G_{>t}|\mathcal{K}_f)}$ , then:

$$\frac{d}{dt}P(\mathcal{K}|G,\mathcal{K}_{mrca},\mathcal{K}_f) = \sum_{i=1}^{n}\sum_{a=1}^{m}\left[\, q^i_{al_i}P(\mathcal{K}_{l_i=a}\mid\mathcal{K}_{mrca},\mathcal{K}_f,G) - q^i_{l_i a}P(\mathcal{K}\mid\mathcal{K}_{mrca},\mathcal{K}_f,G)\right].$$

Therefore, we have shown that migration on a given network, with known types at a root and leaf nodes can be formulated as a continuous time Markov process with generator matrix $Q^i$ for each lineage $i$, with off-diagonal elements given by $q^i_{ab}$. Combined with the lineage independence approximation for $P(\mathcal{K},G)$ presented above, this lets us directly sample lineage types and migration events using the stochastic mapping algorithm described in the main text.

### Supplementary figures and tables

#### Inference for A/H5N1

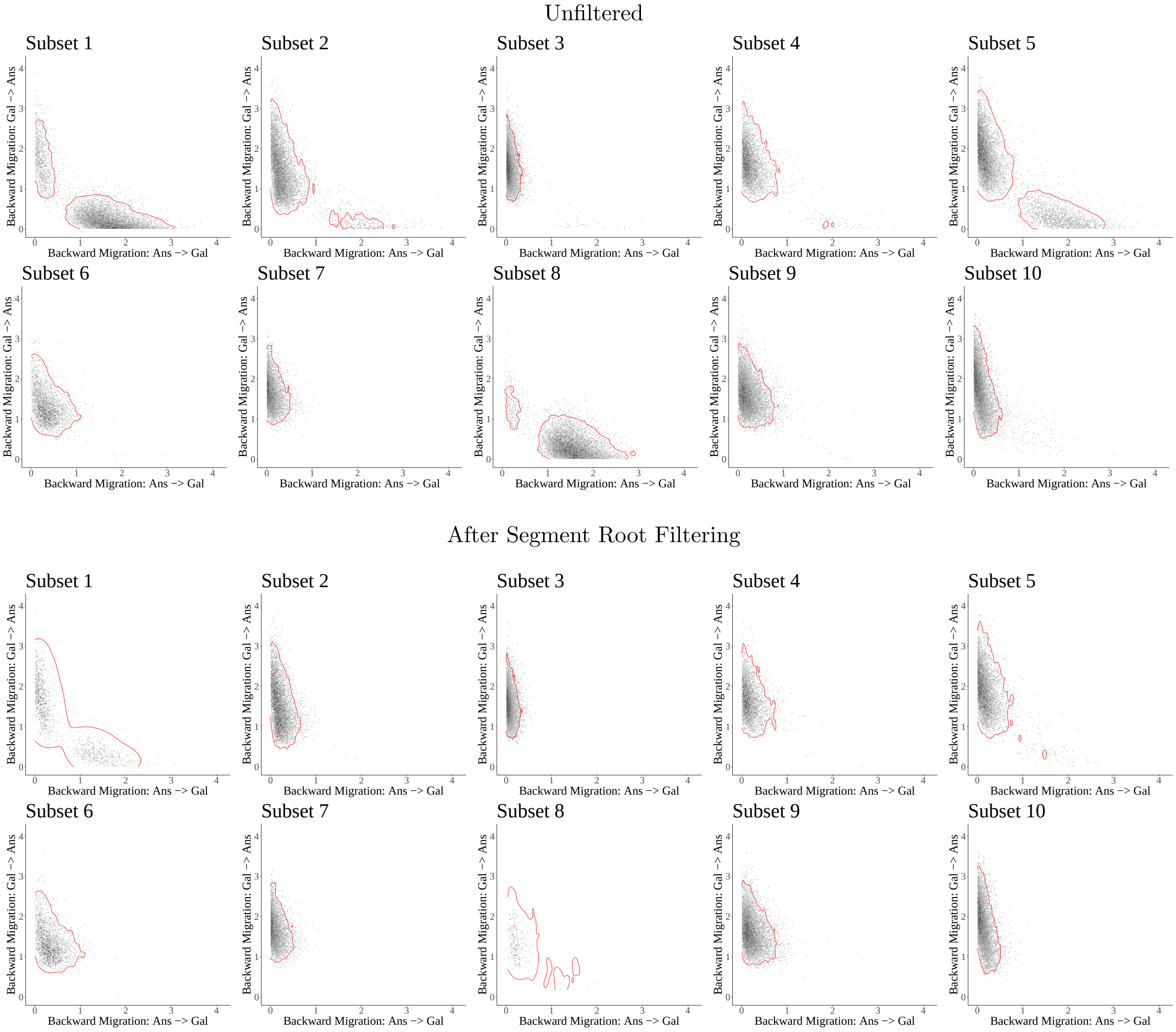


Figure S1: **2D density of backward in time migration rate posterior estimates obtained by SCoRe before and after segment root filtering.** Red contour line marks the 95% HPD area for the estimated 2D density.

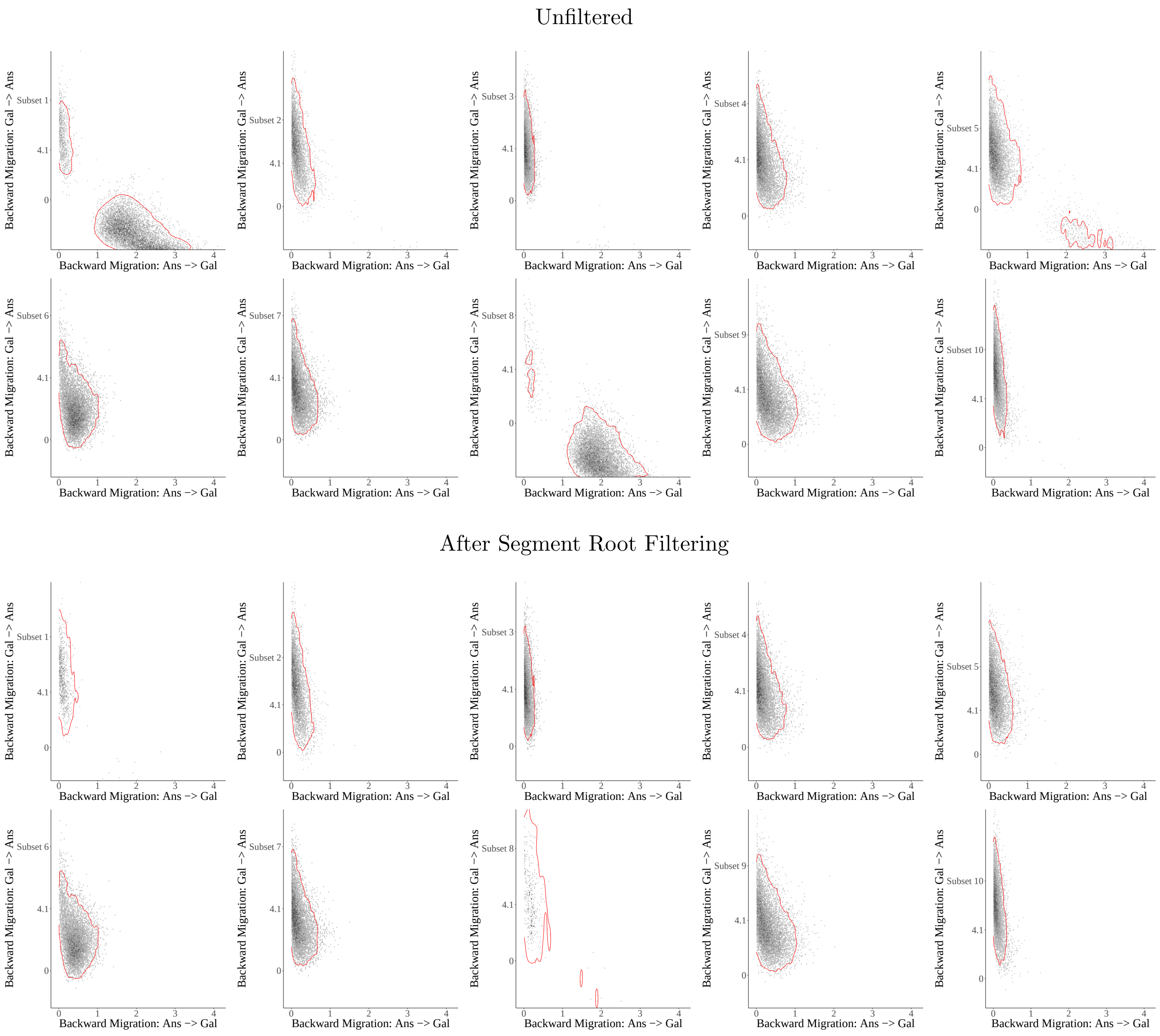


Figure S2: **2D density of backward in time migration rate posterior estimates obtained by MASCOT before and after segment root filtering.** Red contour line marks the 95% HPD area for the estimated 2D density.

#### Unfiltered

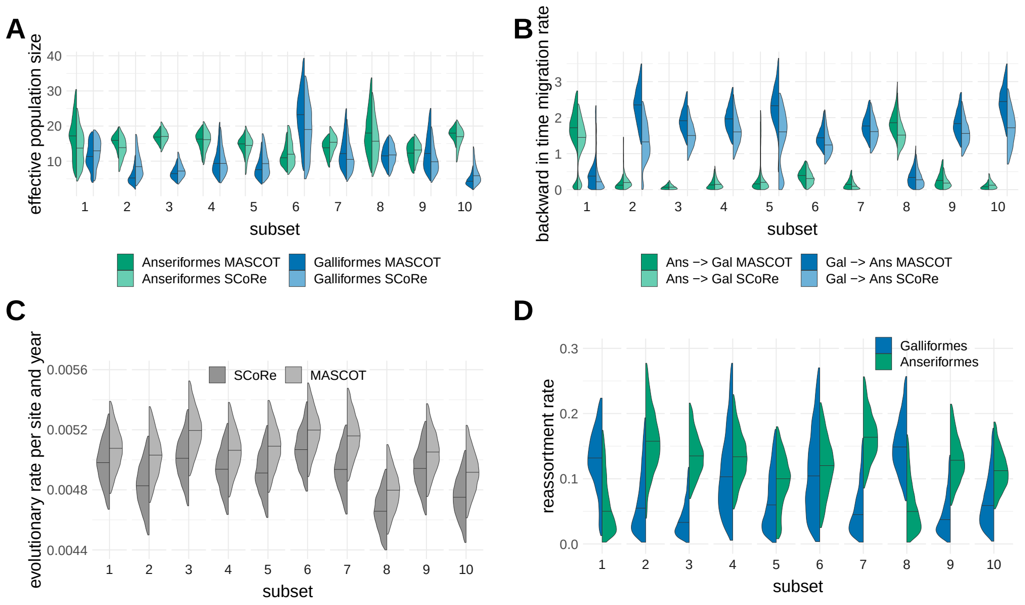


Figure S3: **Posterior distributions of model parameters for all 10 runs with <u>no filtering</u>**. The distributions are compared for both host types (Anseriformes and Galliformes) and for MASCOT and SCoRe packages. For both methods subsets 1,2,5,6 and 8 suffer from uncertainty and bimodality when no filtering is applied. **A** Comparison of effective population sizes. **B** Comparison of inferred backwards in time migration rates for Anseriformes (Ans) and Galliformes (Gal). **C** Comparison of inferred clock rates. SCoRe obtains lower estimates than MASCOT. **D** Comparison of reassortment rates inferred by SCoRe for two bird orders.

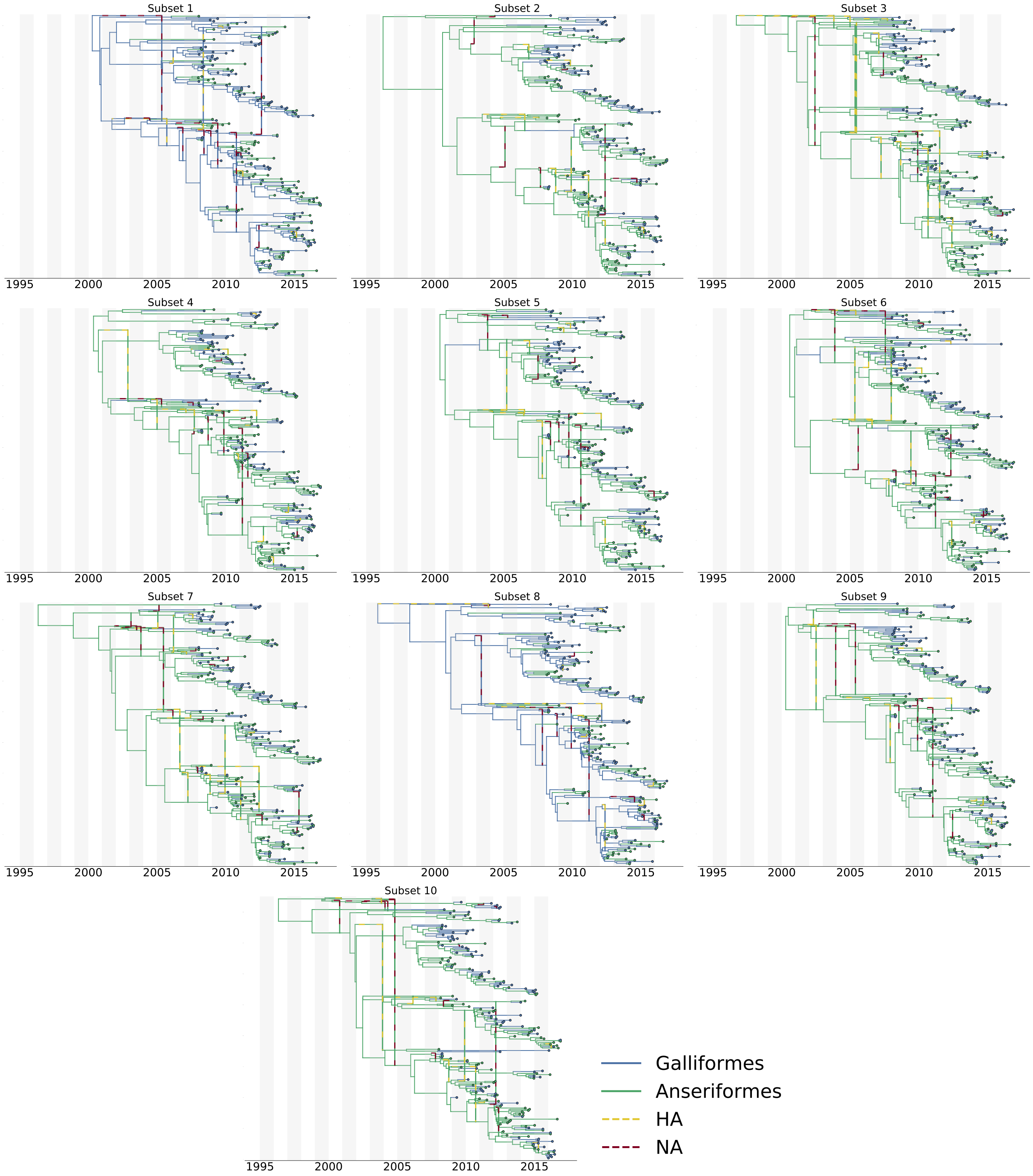


Figure S4: MCC networks of 10 random A/H5N1 subsets, unfiltered. Dashed lines show segment movement at reassortment events. Intermediate migration nodes not displayed.

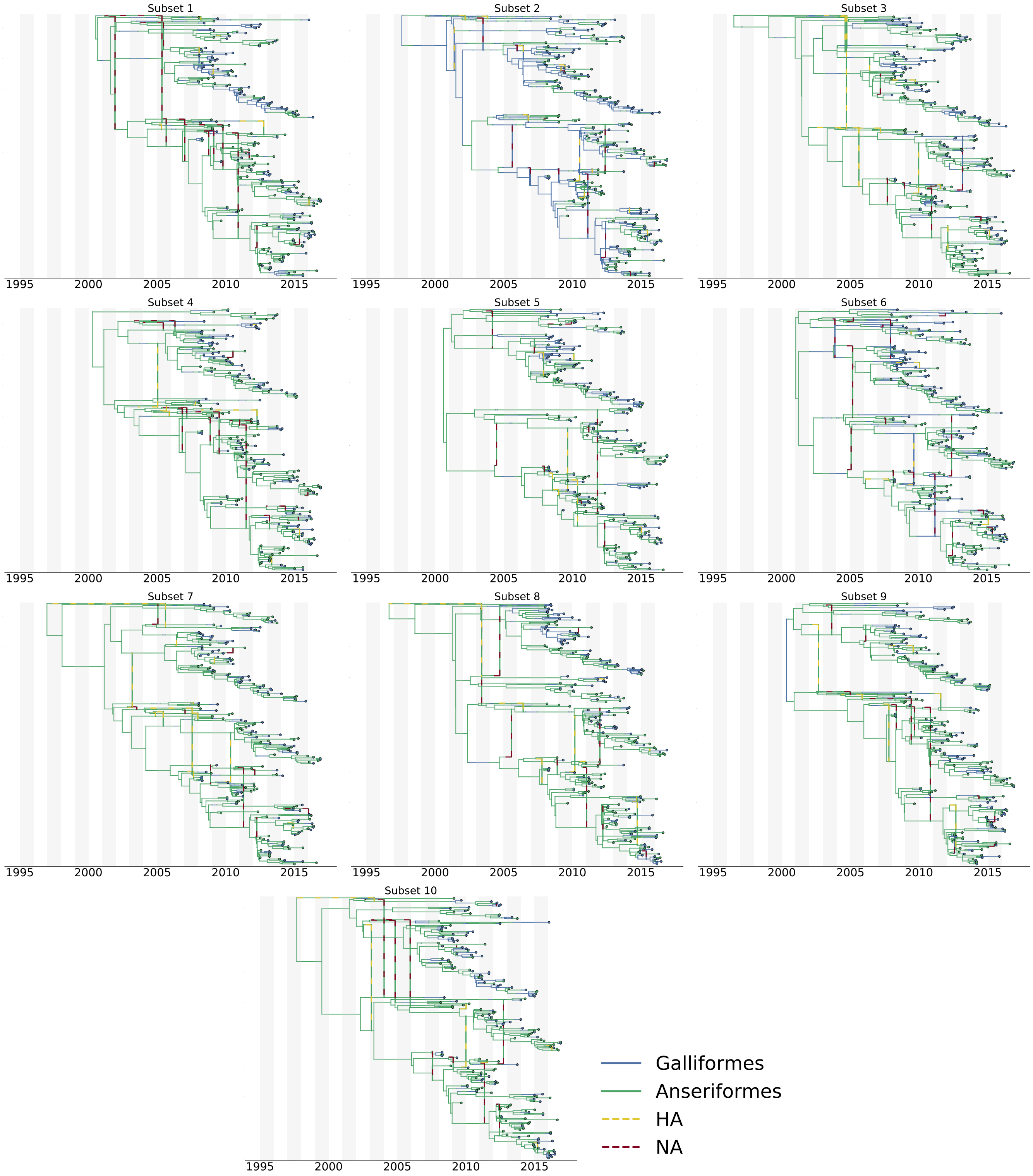


Figure S5: Maximum posterior networks of 10 random A/H5N1 subsets, unfiltered. Dashed lines show segment movement at reassortment events. Intermediate migration nodes displayed.

#### Segment Root Filtering

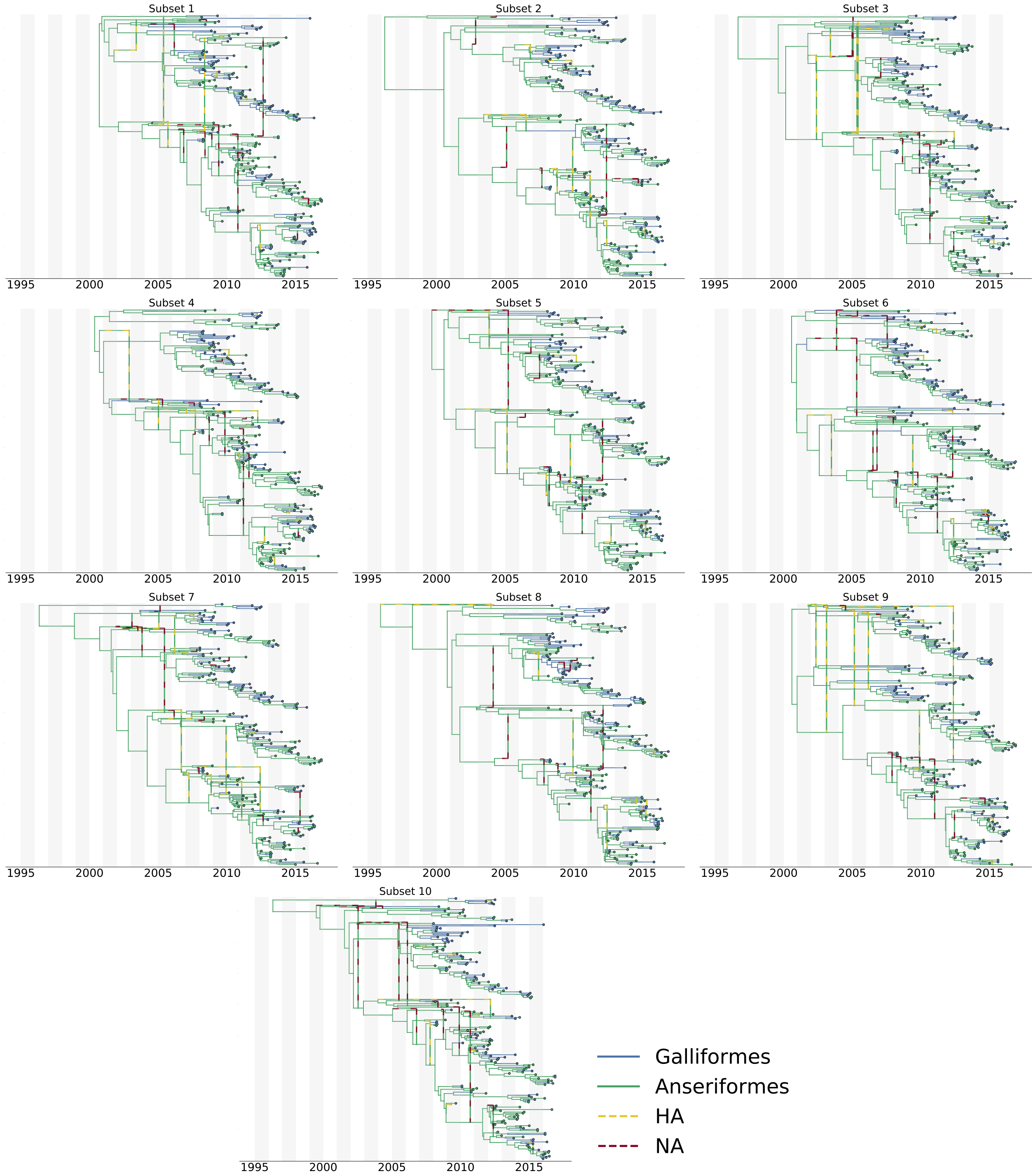


Figure S6: MCC networks of 10 random A/H5N1 subsets, segment root type filtering. Dashed lines show segment movement at reassortment events. Intermediate migration nodes not displayed.

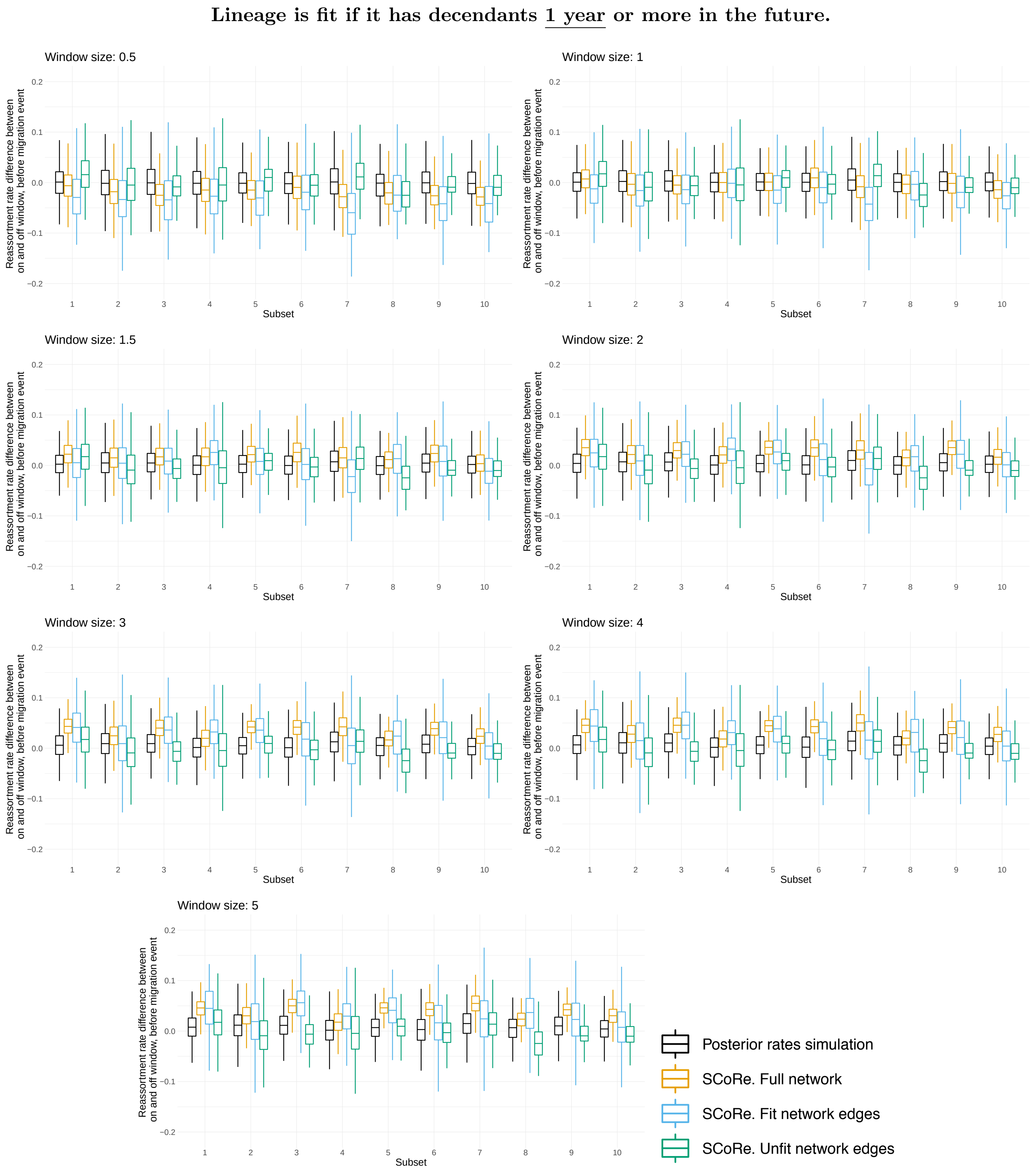


Figure S7: **Reassortment and migration correlation. Difference for posterior "on" and "off window" reassortment rates (y-axis) for 10 random subsets (x-axis).** Each plot corresponds to different sizes (in years) of the window before migration event: 0.5,1.0,1.5,2.0,3.0,4.0 or 5.0. Lineage is fit if it has decendants 1 year or more in the future. For each run, we compare distributions when (a) simulating (no sequencing data) with the posterior rates obtained by the inference or (b) inferring from the sequencing data under the SCoRe model. Since there is a previously defined increase in reassortment due to fitness, which is informed by the data, we show the reassortment rate difference for full network and separately for fit and unfit network edges.

**Lineage is fit if it has decendants 2 years or more in the future.**

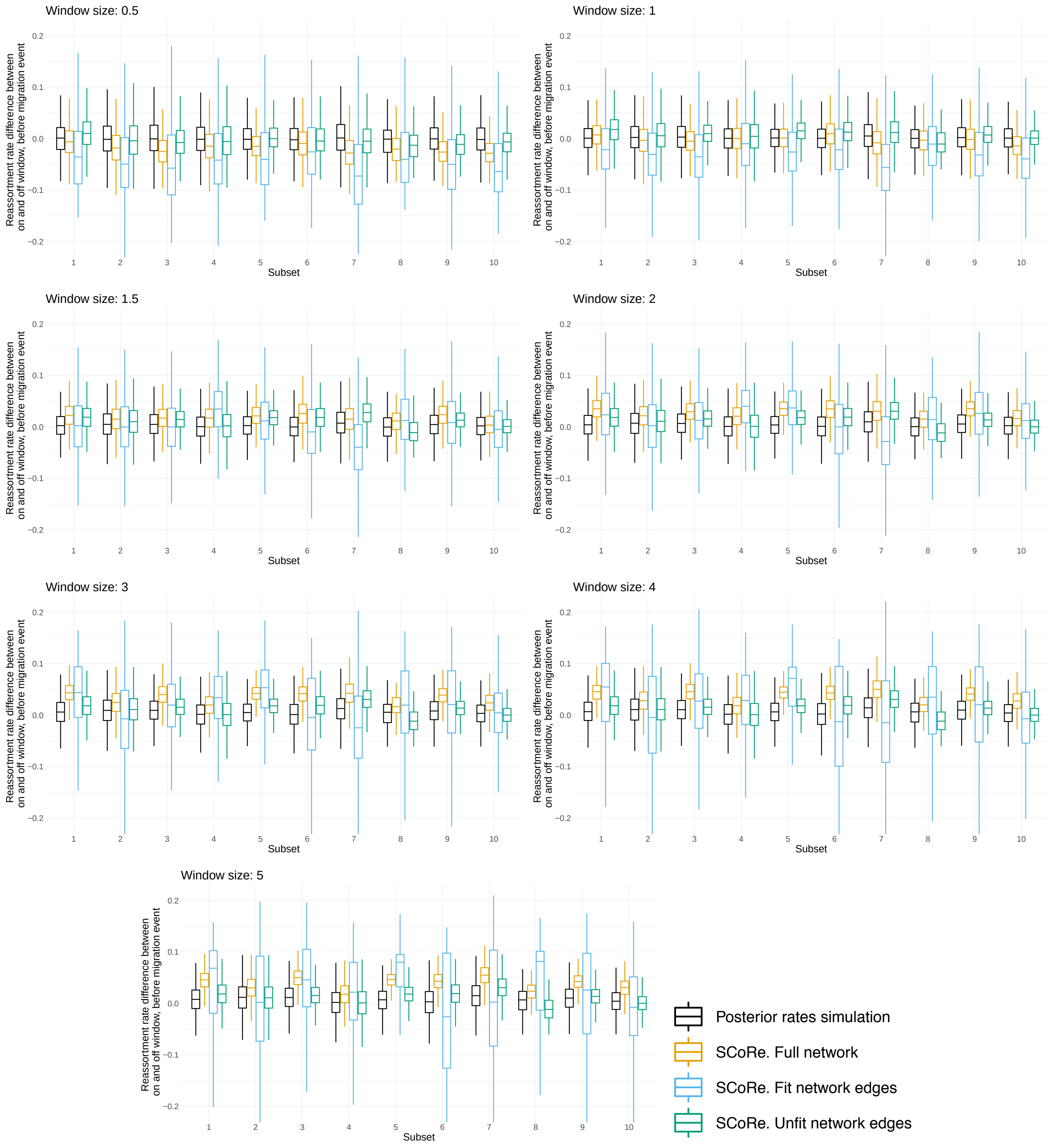


Figure S8: **Reassortment and migration correlation. Difference for posterior "on" and "off window" reassortment rates (y-axis) for 10 random subsets (x-axis).** Each plot corresponds to different sizes (in years) of the window before migration event: 0.5,1.0,1.5,2.0,3.0,4.0 or 5.0. Lineage is fit if it has decendants 2 years or more in the future. For each run, we compare distributions when (a) simulating (no sequencing data) with the posterior rates obtained by the inference or (b) inferring from the sequencing data under the SCoRe model. Since there is a previously defined increase in reassortment due to fitness, which is informed by the data, we show the reassortment rate difference for full network and separately for fit and unfit network edges.

**Lineage is fit if it has decendants 4 years or more in the future.**

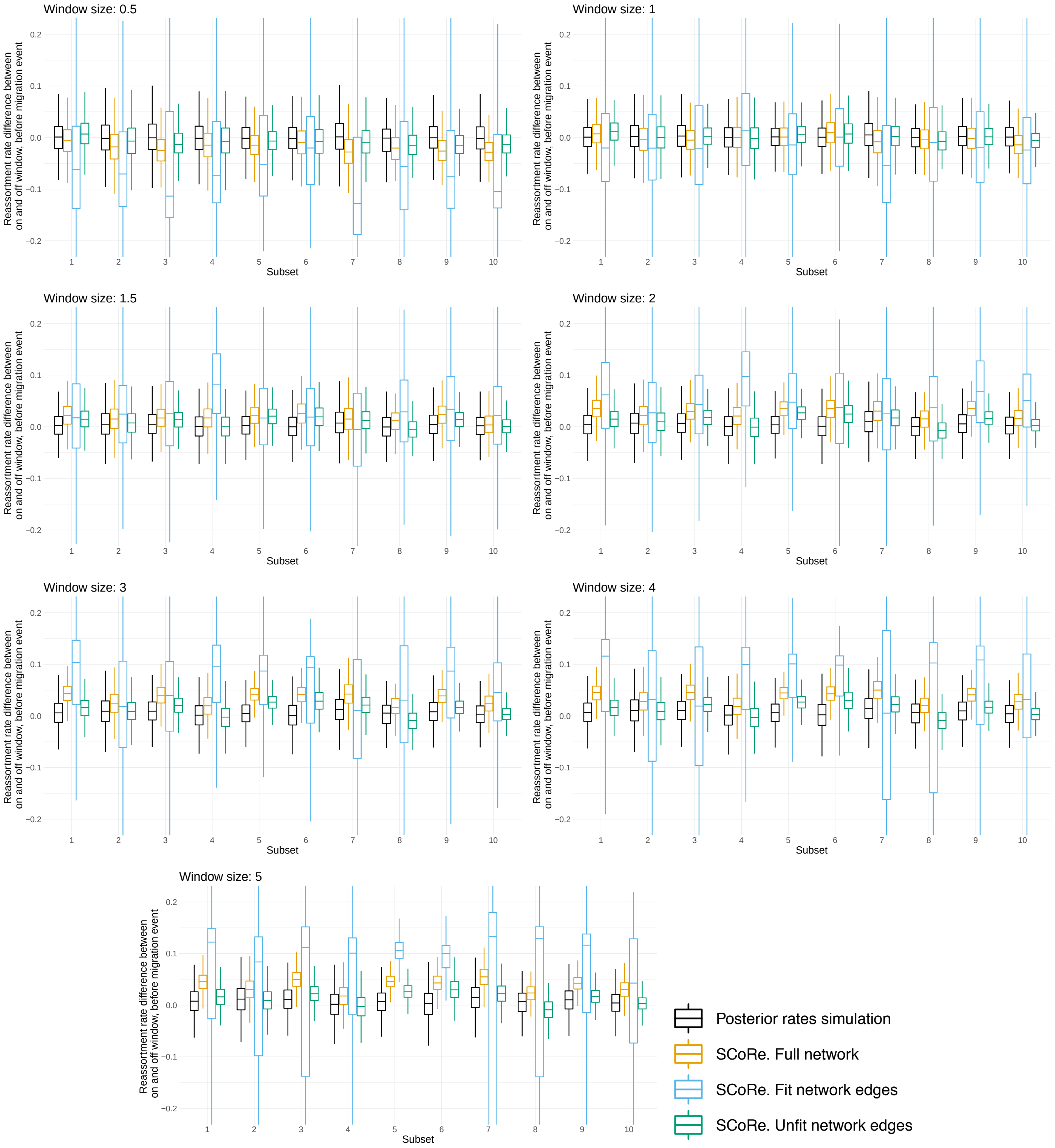


Figure S9: **Reassortment and migration correlation. Difference for posterior "on" and "off window" reassortment rates (y-axis) for 10 random subsets (x-axis).** Each plot corresponds to different sizes (in years) of the window before migration event: 0.5,1.0,1.5,2.0,3.0,4.0 or 5.0. Lineage is fit if it has decendants 4 years or more in the future. For each run, we compare distributions when (a) simulating (no sequencing data) with the posterior rates obtained by the inference or (b) inferring from the sequencing data under the SCoRe model. Since there is a previously defined increase in reassortment due to fitness, which is informed by the data, we show the reassortment rate difference for full network and separately for fit and unfit network edges.

**Lineage is fit if it has decendants 6 years or more in the future.**

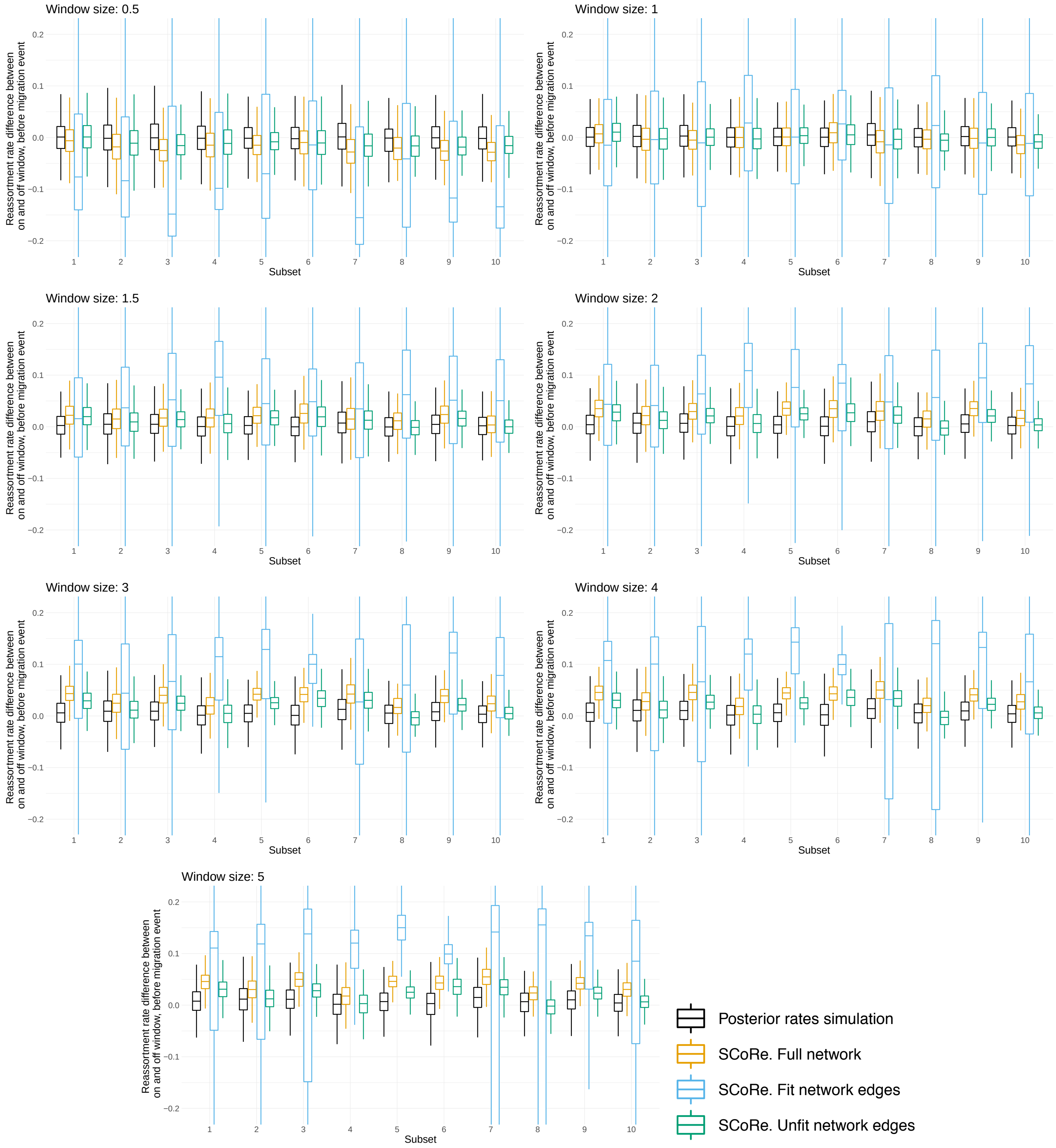


Figure S10: **Reassortment and migration correlation. Difference for posterior "on" and "off window" reassortment rates (y-axis) for 10 random subsets (x-axis).** Each plot corresponds to different sizes (in years) of the window before migration event: 0.5,1.0,1.5,2.0,3.0,4.0 or 5.0. Lineage is fit if it has decendants 6 years or more in the future. For each run, we compare distributions when (a) simulating (no sequencing data) with the posterior rates obtained by the inference or (b) inferring from the sequencing data under the SCoRe model. Since there is a previously defined increase in reassortment due to fitness, which is informed by the data, we show the reassortment rate difference for full network and separately for fit and unfit network edges.

### Validation

#### Implementation validation and true parameter estimation

| Migration prior | Clock rate | Type 1 | | | Type 2 | | |
|---|---|---|---|---|---|---|---|
| | | $N_e$ | $\rho$ | $\mu$ | $N_e$ | $\rho$ | $\mu$ |
| log-normal | high | 93.8 | 91.7 | 90.7 | 96.9 | 93.8 | 97.9 |
| | low | 91.9 | 91.9 | 91.9 | 97.9 | 91.9 | 95.9 |
| | mixed | 90.9 | 92.9 | 92.9 | 97.9 | 91.9 | 95.9 |
| exponential | high | 95.9 | 98.9 | 96.9 | 97.9 | 95.9 | 92.9 |
| | low | 94 | 97 | 94 | 98 | 97 | 93 |
| | mixed | 91.7 | 96.9 | 95.8 | 97.9 | 95.8 | 91.7 |
| Fixed network | | 94.4 | 94 | 93.6 | 95.5 | 94.9 | 94.9 |

Table S1: Percentage of true parameter values falling within 95% HPD interval for simulation study of 2 types. The top six rows shows cases where the network and its parameters were joinly inferred from simulated genetic sequence data for two different migration priors and high ($5 \times 10^{-3}$), low ($5 \times 10^{-4}$) or mixed (2 segments with high and 2 with low) clock rates. The bottom row shows results obtained when the true network is known and only its parameters are inferred. True rates and effective population sizes were asymetric in all cases.

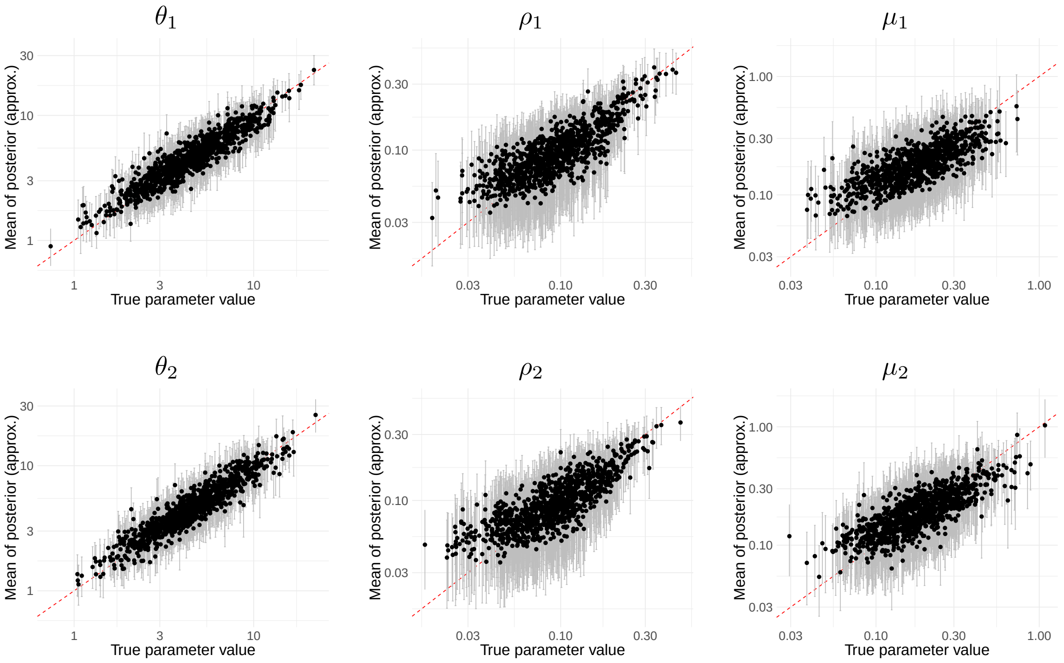


Figure S11: Inference of effective population size ($\theta$), reassortment ($\rho$) and migration ($\mu$) rates from 1000 fixed networks with 4 segments and 2 types. Top row of is for type 1 and bottom – for type 2 parameters. True (x-axis) versus estimated (y-axis) effective population sizes. Grey bars are 95% confidence intervals, red marks the x=y curve.

#### Log-normal migration prior

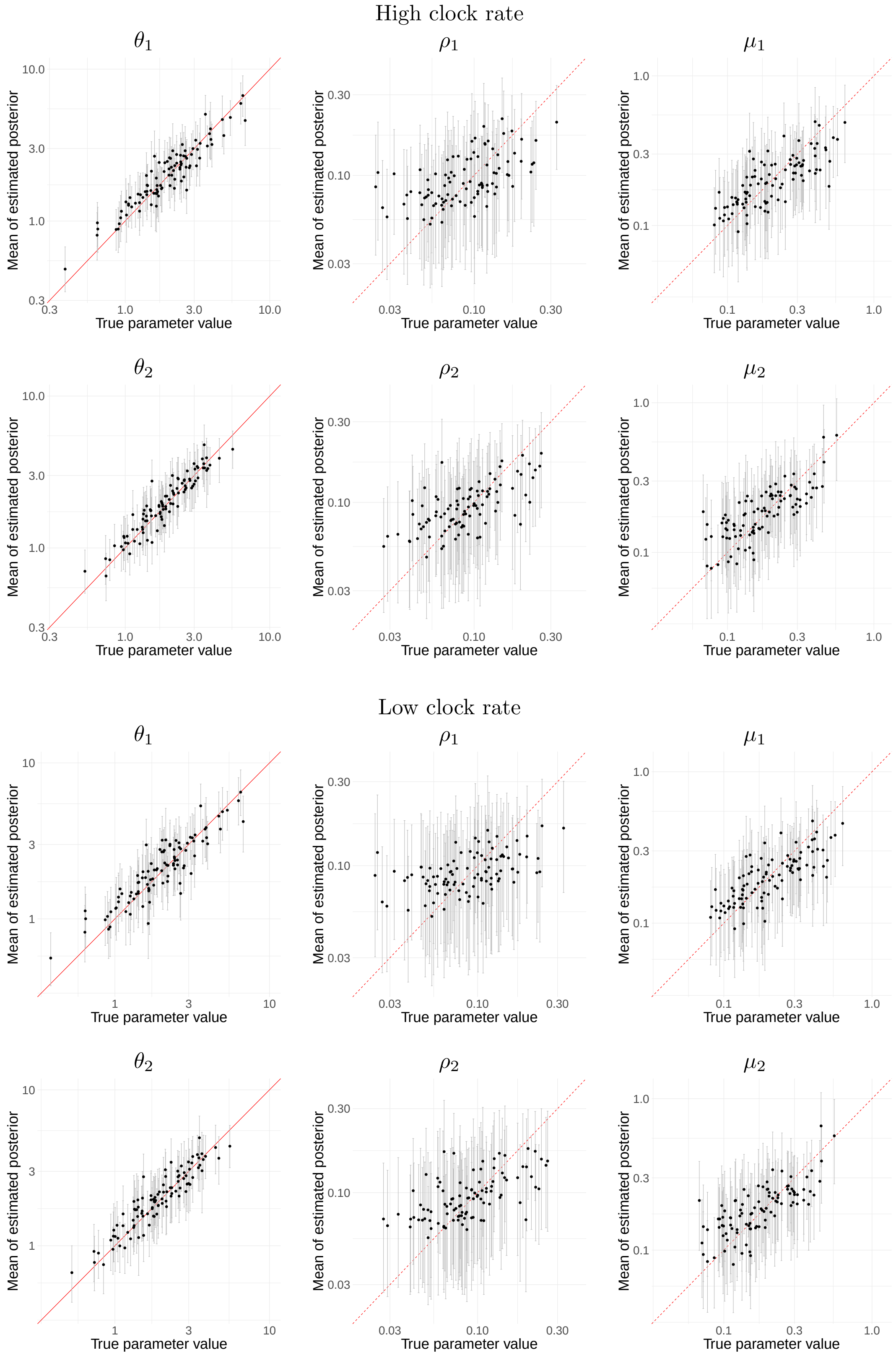


Figure S12: Inference of effective population size ($\theta$), reassortment ($\rho$) and migration ($\mu$) rates from 100 simulated genetic sequence data for 2 types with log-normal migration rate distribution and high ($5 \times 10^{-3}$) or low ($5 \times 10^{-4}$) clock rate for all 4 segments. Top row of is for type 1 and bottom – for type 2 parameters. True (x-axis) versus estimated (y-axis) effective population sizes. Grey bars are 95% confidence intervals, red marks the x=y curve.

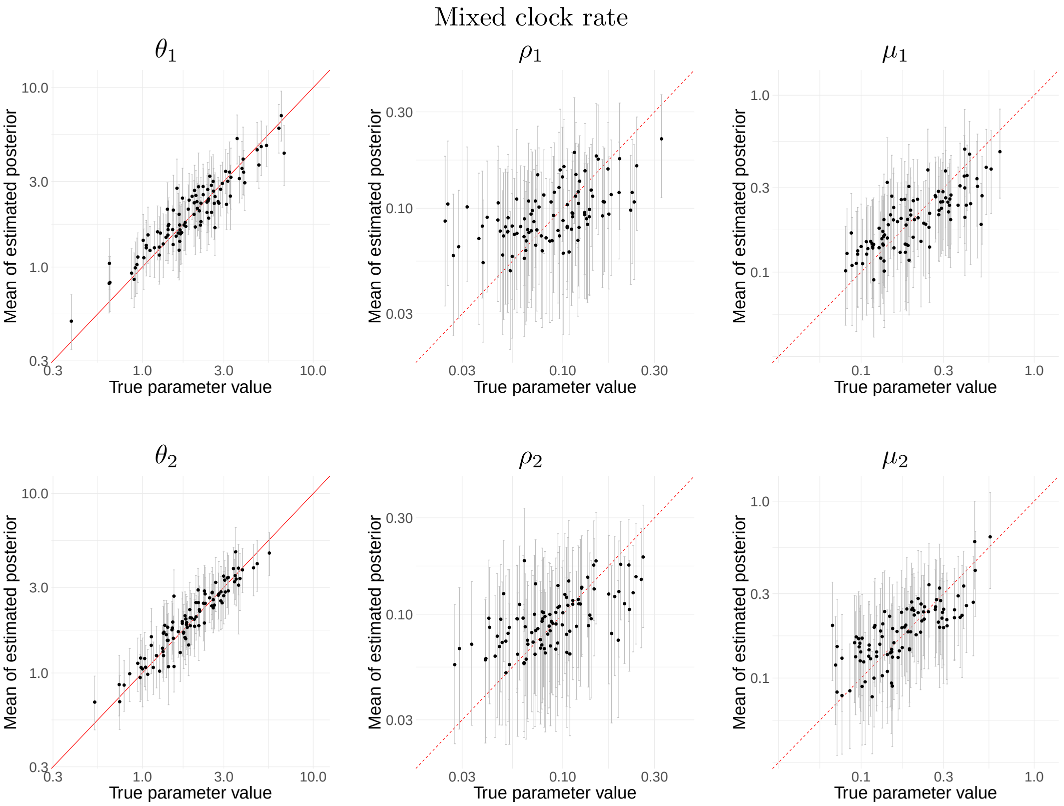


Figure S13: Inference of effective population size ($\theta$), reassortment ($\rho$) and migration ($\mu$) rates from 100 simulated genetic sequence data for 2 types with log-normal migration rate distribution and mixed clock rate ( $5 \times 10^{-3}$ for first 2 segments and $5 \times 10^{-3}$ for remaining 2). Top row of is for type 1 and bottom – for type 2 parameters. True (x-axis) versus estimated (y-axis) effective population sizes. Grey bars are 95% confidence intervals, red marks the x=y curve.

#### Exponential migration prior

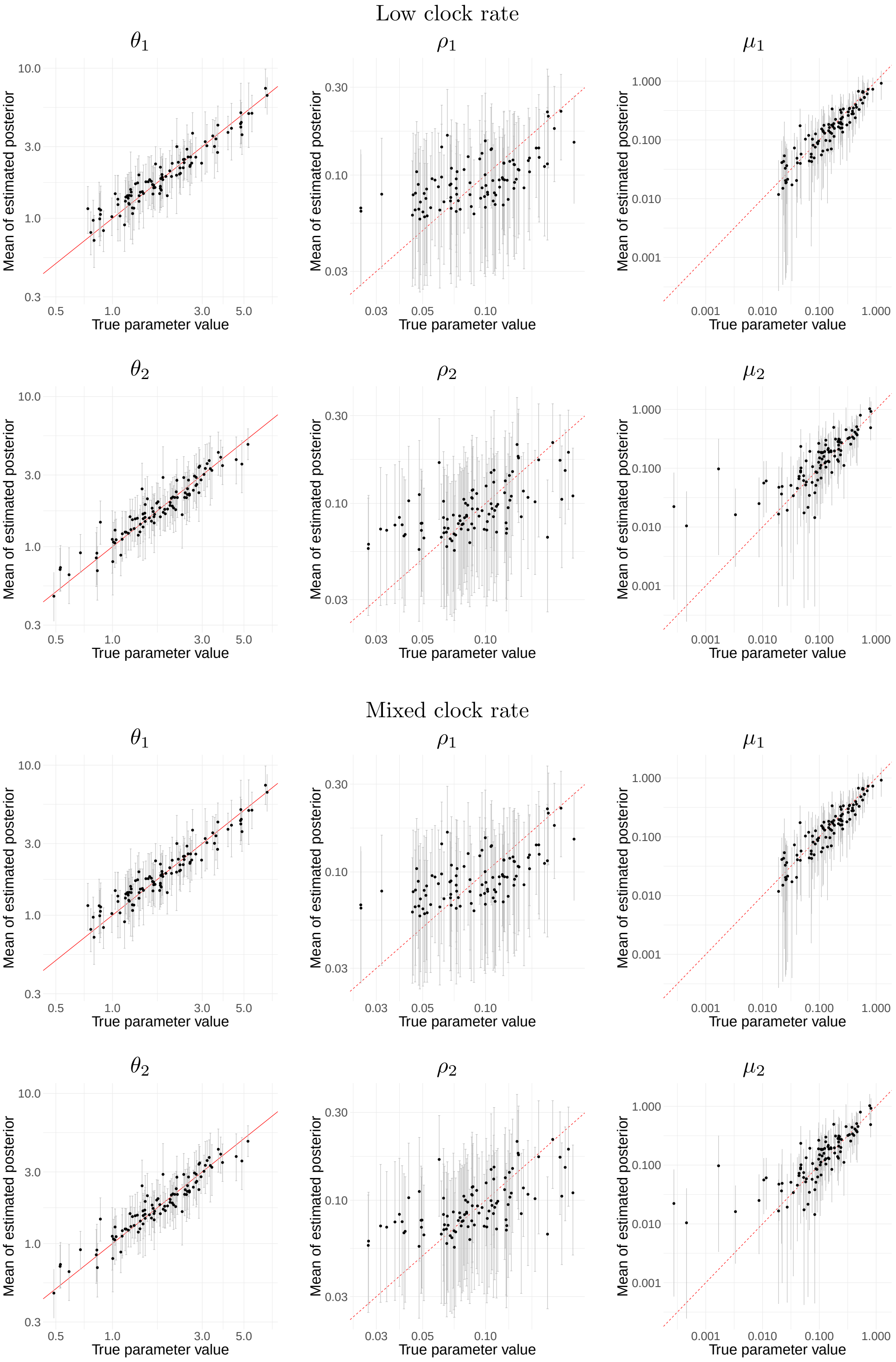


Figure S14: Inference of effective population size ($\theta$), reassortment ($\rho$) and migration ($\mu$) rates from 100 simulated genetic sequence data for 2 types with exponential migration rate distribution and low ($5 \times 10^{-43}$ for all 4 segments) and mixed ( $5 \times 10^{-3}$ for first 2 segments and $5 \times 10^{-3}$ for remaining 2) clock rate. Top row of is for type 1 and bottom – for type 2 parameters. True (x-axis) versus estimated (y-axis) effective population sizes. Grey bars are 95% confidence intervals, red marks the x=y curve.

**Rellative error of ScoRe and MASCOT**

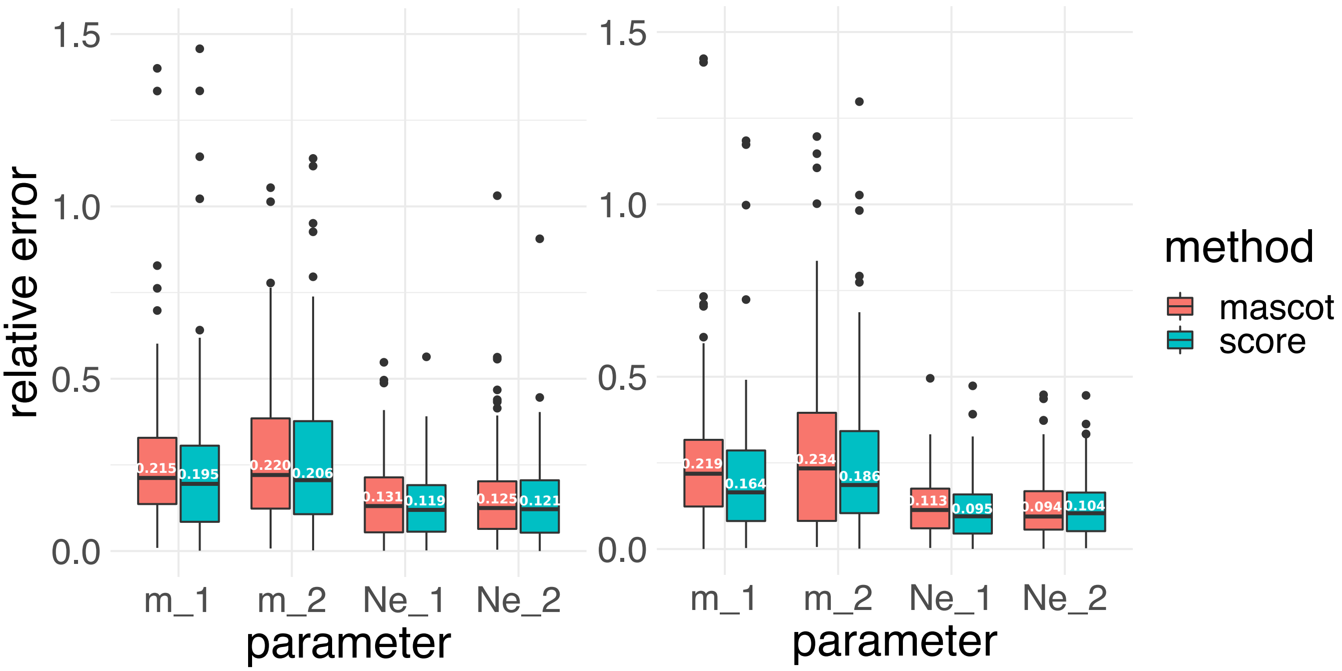


Figure S15: Relative error for parameter inference with SCoRe and MASCOT for low **(right)** and high **(left)** clock rates. Relative error is calculated as described in the main text, for median values of inferred posterior distributions. Horizontal lines mark median of the relative error distribution with its exact value noted in white. Reassortment rate was fixed to true value in SCoRe inference.

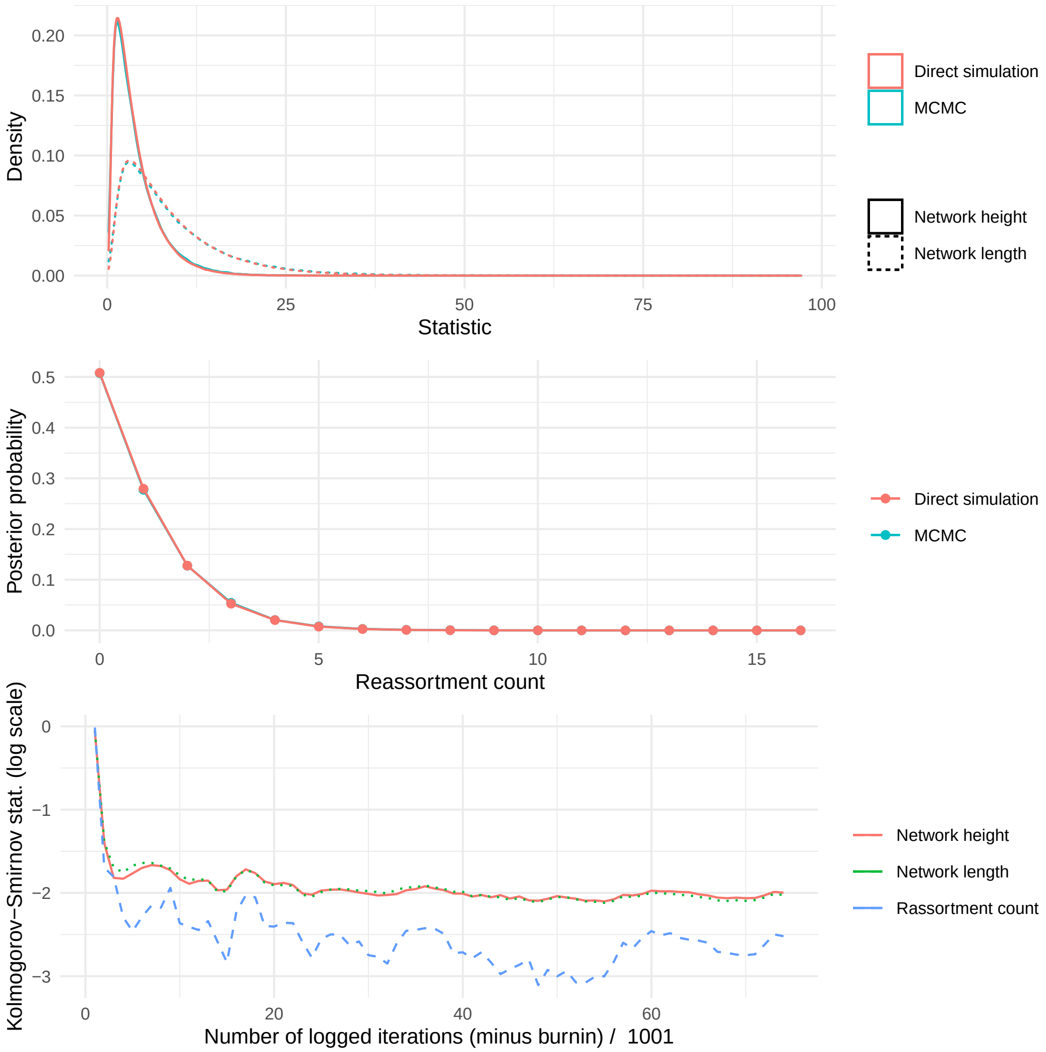


Figure S16: Comparison of network statistics for sampled and simulated structured coalescent with reassortment for 3 taxa of 2 types, each carrying 3 segments. MCMC sampling done by the SCORE-exact. **Top**: Sampled and simulated distributions of network height and length. **Middle**: Sampled and simulated numbers of reassortment events. **Bottom**: Difference between distributions quantified as Kolmogorov Smirnov difference.

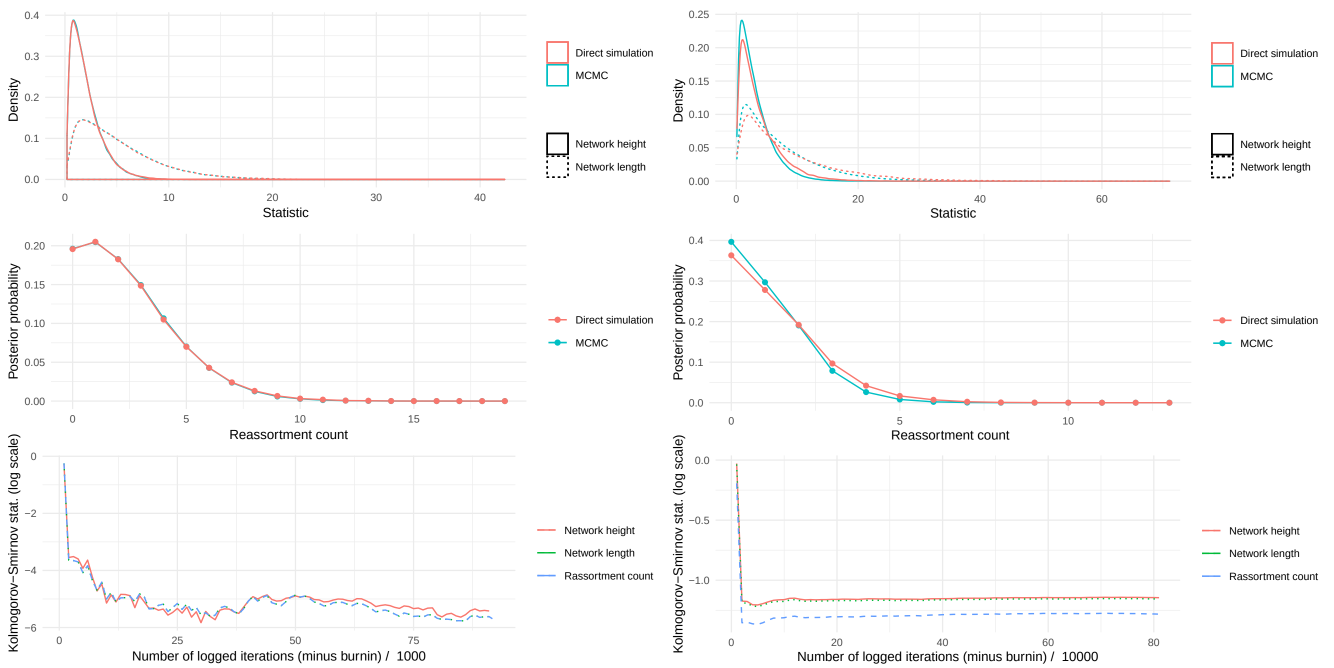


Figure S17: Comparison of network statistics for sampled and simulated coalescent with reassortment (**left**, 3 taxa, each carrying 3 segments) or structured coalescent with reassortment (**right**, 2 taxa of 2 types, each carrying 2 segments). MCMC sampling done by the approximate SCORE variant. **Top**: Sampled and simulated distributions of network height and length. **Middle**: Sampled and simulated numbers of reassortment events. **Bottom**: Difference between distributions quantified as Kolmogorov Smirnov difference.
